# Tuft cells act as regenerative stem cells in the human intestine

**DOI:** 10.1101/2024.03.17.585165

**Authors:** Lulu Huang, Jochem H. Bernink, Amir Giladi, Daniel Krueger, Gijs J.F. van Son, Maarten H. Geurts, Georg Busslinger, Lin Lin, Maurice Zandvliet, Peter J. Peters, Carmen López-Iglesias, Christianne J. Buskens, Willem A. Bemelman, Harry Begthel, Hans Clevers

**Affiliations:** Hubrecht Institute, Royal Netherlands Academy of Arts and Sciences (KNAW) and University Medical Centre Utrecht (UMC) Utrecht, 3584 CT, the Netherlands; Oncode Institute, Hubrecht Institute, 3584 CT Utrecht, the Netherlands; The Princess Maxima Center for Pediatric Oncology, 3584 CS Utrecht, the Netherlands; Department of Clinical Sciences of Companion Animals, Faculty of Veterinary Medicine, Utrecht University, the Netherlands; The Maastricht Multimodal Molecular Imaging Institute, Maastricht University, Maastricht, the Netherlands; Department of Experimental Immunology, Amsterdam Infection & Immunity Institute (AI&II), Amsterdam UMC, University of Amsterdam, Amsterdam, the Netherlands; Pharma, Research and Early Development (pRED) of F. Hoffmann-La Roche Ltd, Basel, Switzerland

**Author notes:** Contributed equally (Lulu Huang, Jochem H. Bernink, Amir Giladi).

## Abstract

In mice, intestinal tuft cells have been described as a long-lived, post-mitotic cell type of which two distinct subsets have been identified, named tuft-1 and tuft-2^1^. By combining analysis of primary human intestinal resection material and intestinal organoids, we identify four distinct human tuft cell states, two of which overlap with their murine counterparts. We show that tuft cell development depends on the presence of Wnt ligands, and that tuft cell numbers rapidly increase upon interleukin (IL)-4 and IL-13 exposure, as reported previously in mouse^2–4^. This occurs through proliferation of pre-existing tuft cells, rather than through increased *de novo* generation from stem cells. Indeed, proliferative tuft cells occur *in vivo* both in fetal and in adult human intestine. Single mature proliferating tuft cells can form organoids that contain all intestinal epithelial cell types. Unlike stem- and progenitor cells, human tuft cells survive irradiation damage and retain the ability to generate all other epithelial cell types. Accordingly, organoids engineered to lack tuft cells fail to recover from radiation-induced damage. Thus, tuft cells represent a damage-induced reserve intestinal stem cell pool in humans.

## INTRODUCTORY PARAGRAPH

The human intestinal tract is responsible for the uptake of food constituents and metabolites, and production of hormones, while adroitly providing protection against pathogens. Its divergent functions are executed by specialized epithelial subsets that are continuously replenished by LGR5^+^ Crypt Base Columnar (CBC) stem cells. Tuft cells represent one such specialized epithelial subset^5^. The majority of our understanding of tuft cell function and ontology derives from studies in mice as there currently exists no *in vitro* approach to generate and manipulate human tuft cells.

In mice, intestinal tuft cells have been described as a long-lived, post-mitotic cell type. Two distinct subsets have been identified, named tuft-1 and tuft-2^1^. Tuft-1 cells express genes that are associated with neuronal interactions, whereas tuft-2 cells appear primarily involved in providing protective immunity against helminths, protozoans, and bacteria^6–9^. Mechanistically, murine tuft cells express the cytokine IL-25, which triggers resident type 2 innate lymphoid cells (ILC2s) to secrete IL-13. This cytokine is thought to signal to the CBC stem cells and to uncommitted progenitor cells to promote the appearance of tuft cells in a positive feedback loop^2–4^. Tuft-2 cells sense microbe-derived metabolic compounds, resulting in enhanced secretion of arachidonic acid-derived lipid mediators, such as prostaglandins and leukotrienes that in turn activate IL-13 producing ILC2s and mucus-secreting goblet cells.

While Lgr5^+^ CBC cells represent the workhorse stem cells of the intestinal epithelium^10^, they are highly susceptible to damage and various mechanisms have been shown to compensate for stem cell loss in mice. One such mechanism involves plasticity of fated crypt progenitors. Thus, dedifferentiation of secretory progenitors^11–14^ and enterocyte progenitors^15^ can restore normal numbers of Lgr5^+^ stem cells. Batlle and colleagues identified a subpopulation of quiescent Lgr5^+^ stem cells marked by Mex3a that can restore stem cell function^16^. Finally, so called revival stem cells (RevSCs), defined by transient induction of Clusterin (Clu) and Sca1, have been proposed to be key to intestinal regeneration, although it is not known from which cells these RevSCs derive^17,18^.

The lack of models to study human tuft cell biology impedes the functional characterization and properties of these cells in detail, while no human intestinal regenerative stem cell has been identified as of yet. Here we use human intestinal organoids in direct comparison with primary intestinal epithelium, to explore human tuft cell development, its subsets and its responses following damage.

## RESULTS

### Wnt signaling drives human intestinal tuft cell development

To generate a model for visualization and tracking of human intestinal tuft cell dynamics and function, we mined published single cell RNA-sequencing resources to select markers that are uniformly, yet specifically, expressed among human tuft cells^19^. We thus identified *AVIL*, a gene encoding for the structural actin-binding protein Advillin, to be highly expressed in tuft cells and not in other intestinal epithelial cells (>192-fold higher expression than any other epithelial type, Fig. 1a), as previously also documented for murine tuft cells^20,21^. Histological examination confirmed the specificity of AVIL expression in primary human small intestinal and colon tissue (Fig. 1b).

**Fig. 1.**
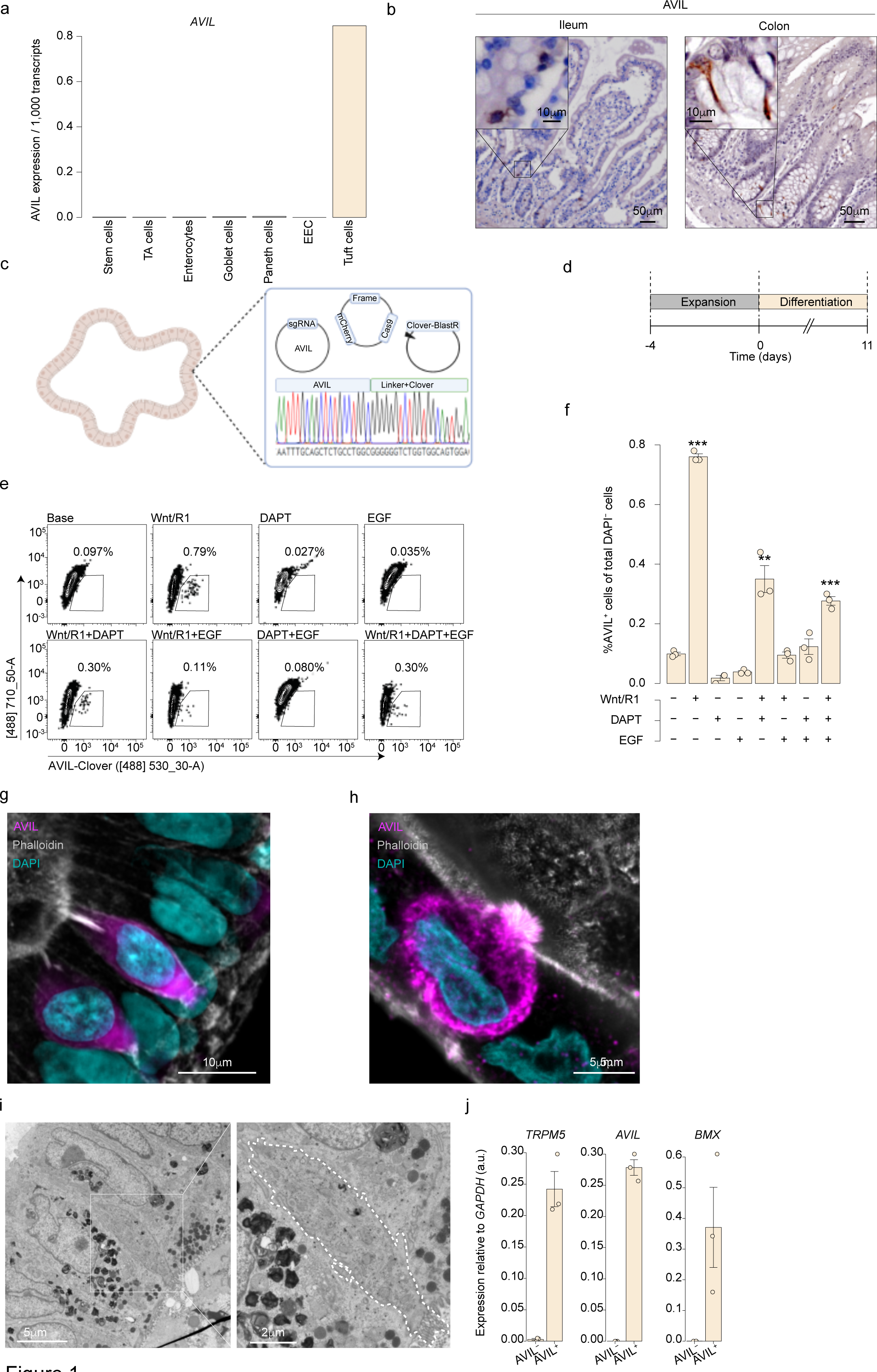
Wnt signaling drives human intestinal tuft cell development. **a,** Mean expression of *AVIL* in a single cell RNA sequencing dataset of human adult small intestinal tissue^18^. n=15,184 single epithelial cells. **b,** Expression of AVIL in histological sections from human adult intestinal tissue. Images are representative of 3 donors with similar results. **c,** Schematics of generating human intestinal organoids with an AVIL-Clover reporter. Organoids were transfected with NHEJ plasmids, single transfected cells were picked and outgrowing clonal organoids were genotyped. n= 4 (donor 1 is shown, see Supplementary Information Fig. 1 for donors 2-4). **d,** Schematics of the experimental set-up for **(e)**–**(f)**. Organoids were cultured in standard human intestinal expansion medium for 4 days, then exposed to various differentiation regimens, in addition to base medium: NAC, B27, and Noggin. **e-f,** Flow cytometric analysis **(e)** and quantification **(f)** of the percentage of AVIL-Clover^+^ cell out of total 4′,6-diamidino-2-phenylindole negative (DAPI^-^) cells cultured in differentiation regimens. Each dot is one well, results are representative of 3 independent experiments on donor 1 (Supplementary Information Fig. 1). Error bars indicate SE. FDR- adjusted one-sided Student’s t-test against the base medium. **g-h,** Fluorescence images of differentiated AVIL-Clover organoids (magenta) counterstained with Phalloidin (F-actin, gray), and DAPI (cyan). n= 2 donors, donor 1 (h) and donor 2 (g). **i,** Transmission electron microscopy (TEM) of tuft cells in differentiated human small intestinal organoids. Donor 2 is shown, also see Supplementary Information Fig. 1. **j,** qPCR quantification of known tuft cell genes in sorted AVIL^+^ and AVIL^-^ cells from AVIL-Clover reporter organoids differentiated for 11 days in human intestinal tuft cell medium (Wnt surrogate, R-spondin1 (CM), Noggin, EGF and DAPT). Each dot is one donor, n= 3 donors (donor 1-2, 4). Error bars indicate SE. TA cells: Transit-Amplifying Cells; EEC: Enteroendocrine cells; R1: R-spondin1 condition medium; SE: standard error. ** P < 0.01, *** P < 0.001.

We generated clonal organoid lines from human duodenum, ileum and colon in which we fused a green fluorescent protein (mClover) to the C-terminus of AVIL using a CRISPR-assisted non-homologous end-joining approach at the *AVIL* locus^22^ (Fig. 1c). Following an expansion phase of 4 days, AVIL-Clover organoids were subjected to multiple differentiation regimens in which growth factors were added either alone or in combination (Fig. 1d). Using flow cytometry for quantification, we observed the highest frequency of AVIL-Clover^+^ cells in the presence of Wnt (Fig. 1e-f, Extended Data Fig. 1a). No higher tuft cell numbers were reached by blocking Notch signaling using the gamma-secretase inhibitor DAPT, and addition of epidermal growth factor (EGF) reduced tuft cell frequency (Fig. 1e-f), although these factors increased overall cell survival (Extended Data Fig. 1b). The specificity of the reporter lines was confirmed by antibody staining for endogenous AVIL (Extended Data Fig. 1c). Like primary tuft cells, organoid-derived AVIL^+^ cells were characterized by flask-shaped contours, and intense bundling of actin fibers towards the tip of the tuft (Fig. 1g-h). Electron microscopy imaging further revealed that, like primary tuft cells, organoid-derived tuft cells formed distinctive lateral protrusions, or interdigitating spinules (Fig. 1i)^23^. In agreement with these observations, purified AVIL^+^ cells expressed high levels of previously described signature tuft cell genes as demonstrated by quantitative polymerase chain reaction (qPCR) (Fig. 1j)^19^.

High Wnt levels are present in the intestinal crypts, and inversely correlate with bone morphogenic protein (BMP) signaling, the latter being a key driver of epithelial differentiation, predominantly active in the intestinal villi^5^. Removal of the BMP inhibitor Noggin and addition of BMP2 and BMP4 to differentiating organoids reduced tuft cell frequency (Extended Data Fig. 1d). The observed dependence on Wnt and on BMP inhibition urged us to explore the cellular localization of tuft cells in the human intestine. Indeed, quantifying tuft cells along the crypt-villus axis in the primary human small intestine tissue revealed that these cells are predominantly restricted to the crypt area (Extended Data Fig. 1e).

### IL-4 signaling drives tuft cell proliferation

Elevated levels of intestinal IL-4 and IL-13, cytokines related to type 2 immunity, drive tuft cell hyperplasia in mice, proposedly by increasing *de novo* generation of tuft cells by Lgr5^+^ stem cells^2–4^. We explored cytokine receptor expression profiles in human intestinal epithelial lineages. Tuft cells (rather than stem cells) expressed the highest level of *IL13RA1*, the gene encoding the shared IL-4 and IL-13 receptor (Fig. 2a). Four days exposure of intestinal organoids to IL-4 and IL-13 (but not to IL-25, IL-27, or SCF (KIT ligand)) induced a ten- to fifteen-fold increase in AVIL^+^ cell frequency, most evident in the absence of EGF (Fig. 2b-c, Extended Data Fig. 2a-b). These results raised the question whether proliferation of *bona fide* tuft cells, rather than differentiation of stem or transit-amplifying (TA) cells to tuft cells, is the main driver of elevated tuft cell numbers following exposure to IL-4 and IL-13. Notably, imaging of organoids revealed a pattern of clustered tuft cells (Extended Data Fig 2c-d), suggestive of self-expansion, while IL-4/IL-13-untreated organoids yielded a more scattered profile (Fig. 2d). Moreover, a substantial fraction of IL-4/IL-13-induced tuft cells stained for KI67, pointing towards local proliferation of differentiated tuft cells (Fig. 2e, Supplementary Video. 1). These findings were not restricted to *in vitro* models, as staining for KI67 and AVIL in primary human small intestinal tissues also sporadically marked double positive cells (Fig. 2f, Extended Data Fig. 2e-f).

**Fig. 2.**
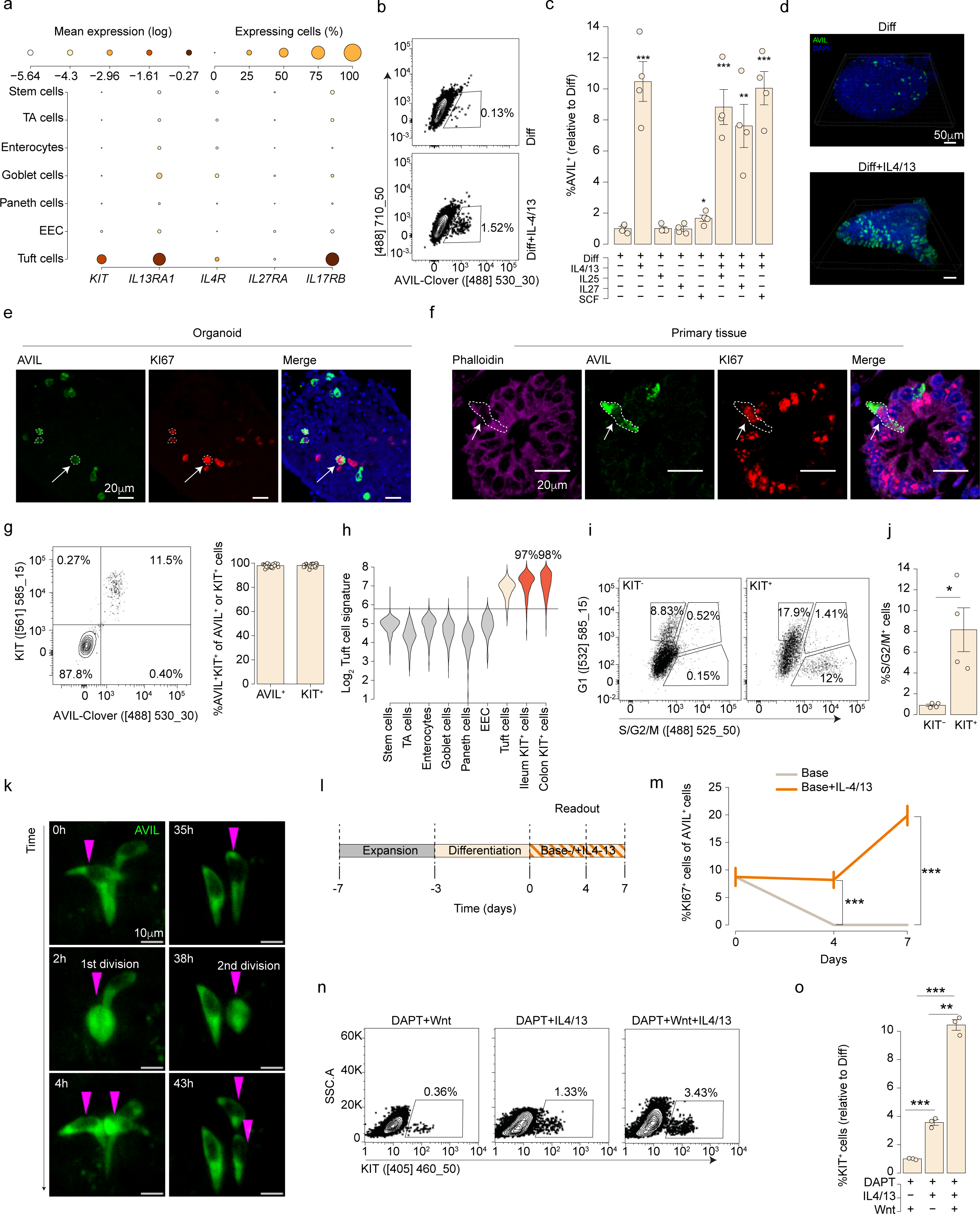
IL-4 signaling drives mitotic tuft cells. **a,** Expression of selected cytokine receptors across the scRNA-seq dataset as in (**1a**). Dot color relates to size-normalized mean expression values and dot size to fraction of expressing cells. **b-c,** Representative flow cytometric analysis **(b)** and quantification **(c)** of AVIL^+^ cell frequency in AVIL-Clover reporter organoids, differentiated for 4 days in tuft cell differentiation medium supplemented with the indicated recombinant proteins. Each dot represents one well. Experiment was performed on 4 donors (donor 1 is shown; for donors 2-4, see Supplementary Information Fig. 2). Error bars indicate SE. FDR-adjusted one-sided Student’s t-test against the Diff medium. **d,** Fluorescence images of AVIL-Clover organoid (green) co-stained with DAPI (blue) differentiated in tuft cell differentiation medium with or without IL-4 and IL-13. n= 3 independent experiments on donor 1. **e,** Fluorescence image of an AVIL-Clover organoid (green) co-stained with KI67 (red) and DAPI (blue). Organoids were differentiated for 4 days in tuft cell medium with IL-4 and IL-13. n= 3 organoid lines derived from 2 donors (one line of donor 1 is shown; for donor 2 see Supplementary Information Fig. 2, for second line of donor 1 see Supplementary Video 1). **f,** Fluorescence image of histological sections of human ileum tissue co-stained for Phalloidin (F-actin, magenta), AVIL (green), KI67 (red) and DAPI (blue). n= 3 donors (Extended Data Fig. 2e). **e-f,** Arrows and dashed lines indicate cells showing AVIL and KI67 overlap. **g,** Representative flow cytometric analysis (left) and quantification (right) of KIT (phycoerythrin, PE) labeled AVIL-Clover organoids cultured in tuft cell differentiation medium supplemented with IL-4 and IL-13. Experiments were performed on 3 donors. Left: donor 1 is shown; for donor 2, 4, see Supplementary Information Fig. 2; right: each dot is one well, results are pooled from 3 donors (donor 1-2, 4). Error bars indicate SE. **h,** Distribution of tuft cell gene expression signature across different epithelial populations from human adult intestine tissue^18^, as well as within sorted populations of KIT^+^ cells from primary ileum and colon. Horizontal line indicates an optimal separation based on the unenriched intestine dataset. Percentage of classified tuft cells in KIT^+^ populations is indicated. The tuft cell signature is based on 222 core tuft cell genes shown in Fig. 3d, f. n=311 ileal KIT^+^ cells, and 271 colon KIT^+^ cells. **i-j,** Representative flow cytometric analysis **(i)** and quantification **(j)** of S/G2/M fraction in KIT^+^ or KIT^-^ cells in ileum FUCCI reporter organoid line. Organoids were differentiated for 3 days in tuft cell differentiation medium with IL-4/IL-13. Each dot is one well. Results are pooled from 3 independent experiments. Error bars indicate SE. Two-sided Student’s t-test. **k,** Snapshots of continuous divisions of AVIL^+^ cells by live-cell imaging. AVIL-Clover reporter organoids were differentiated in tuft cell differentiation medium with IL- 4/IL-13 (Supplementary Video 4). n= 2 donors (donor 1 is shown; for donor 2, see Extended Data Fig. 3c, Supplementary Video 5). **l,** Schematics of the experimental set-up for **(m)**. Organoids were differentiated for 3 days in tuft cell differentiation medium, then medium was switched to either with or without IL-4/13 in base medium (NAC, B27, Noggin), frequency of KI67^+^AVIL^+^ co-stained cells was determined by staining. **m,** Percentage of KI67^+^ cell of AVIL^+^ cells on day 0, 4 and 7. Results are pooled from 3 independent experiments on two organoid lines, > 600 AVIL^+^ cells per time point were quantified (donor 1, see Supplementary Information Fig. 2). Error bars indicate SE. Two-sided Student’s t-test. **n-o,** Representative flow cytometric analysis (**n**) and relative quantification (**o**) of the percentage of KIT^+^ cells differentiated for 3 days in DAPT, with or without addition of Wnt and IL-4/IL-13. Each dot is one well. Results are pooled from two experiments on FUCCI lines (donor 1). Experiment was repeated on AVIL-Clover lines from donors 1,3, see Supplementary Information Fig. 2. Error bars indicate SE. Two-sided Student’s t-test. Diff: human tuft cell differentiation medium; TA: Transit-Amplifying Cells; EEC: Enteroendocrine cells; SE: standard error. * P < 0.05, ** P < 0.01, *** P < 0.001.

To further substantiate these findings, we integrated the Fluorescent Ubiquitination Cell Cycle Indicator (FUCCI) in small intestine organoids using a transposon-based system, allowing to track and quantify the mitotic state in organoids. We identified KIT as a tuft cell-specific surface marker (Fig. 2a, Extended Data Fig. 2g). Indeed, KIT staining displayed a >98% overlap in multiple AVIL-Clover reporter organoid lines (Fig. 2g). This overlap was maintained following exposure to various stimuli (Extended Data Fig. 2h-i), confirming KIT as a robust antigen that specifically marks tuft cells among human intestinal epithelial cells. We further confirmed KIT specificity in both healthy and inflamed intestines, by analyzing publicly available scRNA-seq data of human colon epithelium, obtained from a cohort of pediatric/adult healthy and IBD tissues^24^ (Extended Data Fig. 2j). Likewise, scRNA-seq of KIT-sorted cells from human adult ileum and colon primary tissue revealed that >97% of sorted cells exhibited a tuft cell transcriptional signature (Fig. 2h, Extended Data Fig. 2k).

Thus, KIT specificity allowed us to simultaneously label tuft cells and track their cell cycle progression with the FUCCI construct by flow cytometry analysis. Following IL-4 and IL-13 stimulation, up to 8% of the KIT^+^ fraction entered the S/G2/M-phase, while the KIT^-^ fraction remained quiescent (< 1% cycling) (Fig. 2i-j). Of note, non-cytokine exposed tuft cells were occasionally found in the S/G2/M-phase (Extended Data Fig. 3a, Supplementary Video 2). Using live-cell imaging, we observed IL-4/IL-13-stimulated AVIL^+^ cells which divided two times over a 36-hour period, corroborating their capacity to enter a mitotic state (Fig. 2k, Extended Data Fig. 3b-c, Supplementary Video 3-5). In agreement with these findings, IL-4 and IL-13 stimulation promoted a continuous phase of mitotic tuft cell as determined by FUCCI organoids (Supplementary Information Fig. 2) and a steady increase in KI67^+^ AVIL^+^ cell frequency over a 7-day period (Fig. 2l-m). These processes were enhanced in the presence of Wnt signaling (Fig. 2n-o). Importantly, exposure of IL-4 and IL-13 to organoids that did not contain tuft cells (i.e., organoids differentiated in the absence of Wnt), did not induce proliferative tuft cells(Extended Data Fig. 3d). Neither did we observed proliferative tuft cells in sorted and cultured stem cells in expansion medium in the presence of IL-4 and IL-13 over a 7-day period (Extended Data Fig. 3e). This uncoupled IL-4/IL-13-induced tuft cell proliferation from induced differentiation of precursor cells. Taken together, we found tuft cells to be capable of robust proliferation following type-2 cytokine stimulation *in situ*.

### IL-4 signaling shifts the balance in tuft cell subtypes/states

To gain better understanding of tuft cell function and heterogeneity, we obtained single-cell RNA-sequencing data from 953 ileum organoid-derived epithelial cells, grown in tuft cell differentiation medium with and without exposure to IL-4/IL-13. We used the AVIL-Clover signal to select for tuft cells, resulting in 580 high quality AVIL-expressing cells (Extended Data Fig. 4a-b), and 373 cells from other epithelial lineages (Fig. 3a, Extended Data Fig. 4c). The Metacell package^25^ grouped cells into transcriptionally homogeneous subsets, identifying four distinct clusters, which we succinctly named tuft-1 to -4 (Fig. 3b-c). Non-tuft cells were identified as *LGR5*^+^ stem cells and two types of goblet cells (Fig. 3c, Extended Data Fig. 4c). All four tuft cell clusters shared a unique tuft cell core gene expression program (absent from the non-tuft cell fraction), including the genes *ALOX5AP*, *POU2F3*, *LRMP*, *GNG13*, *BMX, AVIL, and KIT* (Fig. 3d, Extended Data Fig. 4d). We also identified some genes not previously linked to tuft cells, such as the neuropeptide Neuromedin U (*NMU*), and the tyrosine-protein kinase *HCK*. Relative tuft cell subset distribution largely depended on medium composition: addition of IL-4 and IL-13 caused a shift towards tuft-3 and tuft-4 (comprising 14.25% and 79% of all IL-4/13 stimulated tuft cells, respectively), whereas under “homeostatic” differentiation conditions these populations were much rarer (3.5% and 3.25%, Fig. 3e). Note that the ratio of tuft cell substates was determined within treatment groups, and that the absolute number of tuft cells within IL-4/IL-13 treated organoids was approximately 10- to 20-fold higher than within non-cytokine treated organoids (Fig. 2c). Direct comparison of the individual tuft cell substates to other epithelial cells showed unique and shared gene expression patterns for each subset with tuft-3 expressing the largest number (>250) of unique differentially expressed genes (DEG), followed by tuft-2 and tuft-4 (Fig. 3f).

**Fig. 3.**
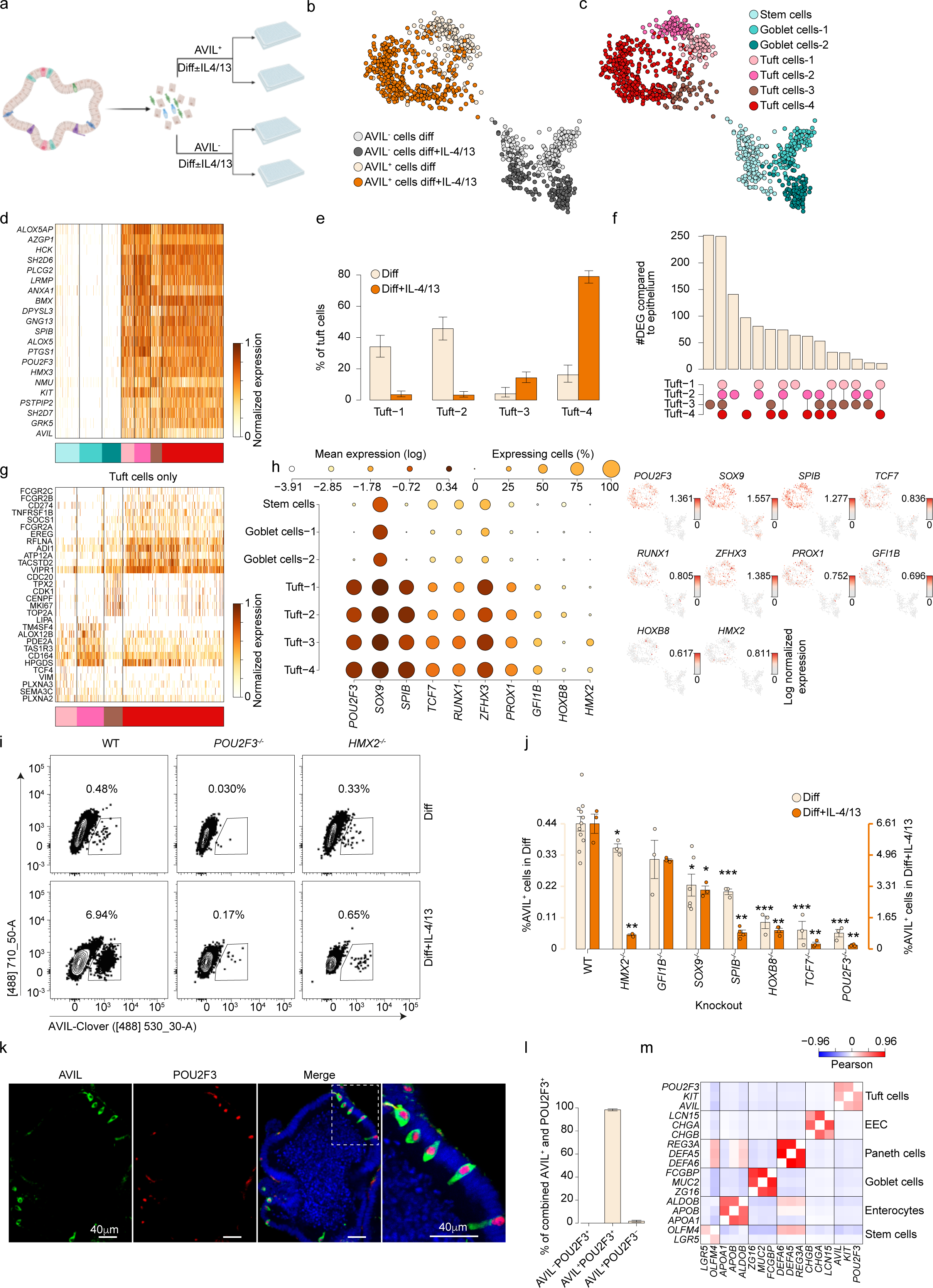
IL-4 signaling shifts the balance in tuft cell states. **a,** Schematic overview of the single cell RNA-seq experiment. Single cells derived from human ileum AVIL-Clover reporter organoids were differentiated in tuft cells medium, with or without IL-4/IL-13 treatment, and sorted based on AVIL expression. **b-c,** Metacell 2D- projection of 953 ileum organoid-derived single cells. Cells are colored by their gating and medium condition **(b)** or by annotation to cell subsets **(c)**. **d,** Gene expression profiles of tuft- cell specific core genes shared by all tuft cell clusters across epithelial subsets. **e,** Distribution of the four distinct tuft cell states in ileum-derived organoids with or without IL- 4/IL-13 stimulation. Error bars indicate 95% confidence in binomial estimation of the mean. **f,** Summary of number of up-regulated differentially expressed genes across the tuft-1-4 states, when compared to non-tuft epithelium. **g,** Gene expression profiles of tuft cell state- specific genes. **h,** Expression of selected genes encoding for transcription factors across ileum organoid-derived epithelial subsets. Left: Dot color relates to mean expression values and dot size to fraction of expressing cells; right: dot color indicates log normalized expression. **b-d, h,** n=953 single cell; **e, g**, 573 single tuft cells. **i-j,** Representative flow cytometric analysis **(i)** and quantification **(j)** of AVIL^+^ cell frequency in organoid mutant lines of selected transcription factors (homozygous knock out). Organoids were differentiated in tuft cell differentiation medium for 11 days or 6 days with IL-4/13 addition. Each dot is one well. Results are representative of 3 independent experiments on donor 1 (Supplementary Information Fig. 3). Error bars indicate SE. FDR-adjusted two-sided Student’s t-test against the WT levels. **k-l,** Representative fluorescence image **(k)** and quantification **(l)** of an AVIL- Clover organoid (green), co-stained for POU2F3 (red) and DAPI (blue). n= 3 donors (donor 1-2, 4). **k**, donor 2 is shown, for donors 1,2 see Extended Data Fig. 5d; **l**, pooled from 3 donors, 993 positive cells from17 organoids were quantified (≥ 5 organoids per donor). Error bars indicate SE. **m**, Gene-pairwise Pearson correlation between markers of the main epithelial types across cells from primary intestinal tissue^18^. Diff: human tuft cell differentiation medium; WT: wildtype; EEC: Enteroendocrine cells; SE: standard error. * P < 0.05, ** P < 0.01, *** P < 0.001.

Gene ontology analysis of the DEG within each tuft cell cluster highlighted the diversity between tuft-1 to -4 (Extended Data Fig. 4e). For example, genes that were enriched in tuft-1 cells are involved in processes of neuron differentiation (e.g., *VIM*, *SEMA3C*), tuft-2 genes are involved in immuno-potent lipid biosynthetic processes such as the synthesis of lipoxins, prostaglandins, and cysteinyl-leukotrienes. Tuft-3 is enriched for genes connected to a mitotic phenotype, with high gene expression levels of *MKI67*, *TOP2A*, *TUBB*, *CKS1B*, and tuft-4 cells express genes that are involved in regulating immune responses and stimulating regeneration, such as genes encoding the SOCS proteins, PD-L1 (*CD274*), *EREG*, *HB-EGF*, *TACSTD2*, amongst others (Fig. 3g). Staining AVIL-Clover reporter organoids with tuft cell-state specific markers further confirmed the observed tuft cell heterogeneity (Extended Data Fig. 4f).

We next asked whether the heterogeneity we observed in organoid tuft cells informed on tuft cell heterogeneity *in vivo*. In our dataset of KIT-sorted tuft cells from human adult ileum and colon primary tissue, we found a small subset of cells exhibiting a large repertoire of tuft-3 genes, implying that tuft cells enter the cell cycle *in vivo* (Extended Data Fig. 4g-h). Combining two published datasets spanning 14 healthy donors^19,26^, we estimated that 3.9% of intestinal tuft cells are cycling in steady state (Extended Data Fig. 4i). We also detected coordinated expression of tuft-4 genes in a gradient-like manner in primary ileum and colon tuft cells, suggesting variable levels of IL-4/13 activation in the homeostatic intestine (Extended Data Fig. 4j-k). We noted that tuft-3 cells *in vivo* exhibit a significantly higher tuft-4 activation signature compared to other tuft cells, linking tuft cell proliferation to cytokine stimulation, similar to our observations *in vitro* (Extended Data Fig. 4l-m).

Taken together, we observed heterogeneity within the human intestinal tuft cell lineage. At steady state, the predominant subsets represent tuft-1 and tuft-2, which phenocopied the two previously described murine intestinal crypt tuft cell subsets^1^. IL-4 signaling triggered induction of cycling tuft-3 cells and the dramatic expansion of tuft-4, the latter not recognized in mouse and likely involved in immune regulation and/or epithelial renewal.

Next, we explored the transcription factor repertoire of tuft cells, identifying *POU2F3*, amongst others, to be uniquely and uniformly expressed by tuft cells in intestinal organoids and tissue (Fig. 3h, Extended Data Fig. 5a). Using conventional CRISPR-Cas9 and base-editing techniques^27^, we generated knockouts of prominent tuft cell-expressing transcription factors in *AVIL*-Clover reporter organoids (Extended Data Fig. 5b) and tested their effect on tuft cell frequency by flow cytometry (Fig. 3i). By quantifying tuft cell frequency in “homeostatic” differentiation medium and IL-4/IL-13-induced medium, we observed that *POU2F3* and the Wnt-signal transducer *TCF7* are essential for tuft cell development (Fig. 3j, Extended Data Fig. 5c). Accordingly, POU2F3 colocalized with AVIL in sections of organoids generated from small and large intestines (Fig. 3k-l, Extended Data Fig. 5d), as well as in a transcriptomic dataset derived from primary intestinal tissue (Fig. 3m). Knocking out *SPIB*, a transcription factor that is shared with M-cells and BEST4^+^ cells^19,28^, partially reduced tuft cell numbers. In addition, we found that organoids mutant for *HMX2* retained normal tuft cell numbers when cultured in “homeostatic” differentiation medium, but essentially lost the tuft cell proliferation response to IL-4 and IL-13, consistent with the differential expression of *HMX2* in tuft-3 and tuft-4 (Fig. 3h). Of note, several conflicting studies in mice reported tuft cell dependence on the secretory lineage-defining transcription factor *ATOH1*^29^. We did not detect *ATOH1* expression in human tuft cells, and ectopic over-expression of *ATOH1* in AVIL-Clover reporter organoids had a negative effect on tuft cell frequency (Extended Data Fig. 5e-f). Of note, *ATOH1* over- expression induced goblet cells and Paneth cells –but not enteroendocrine cells (Extended Data Fig. 5g). Paradoxically, knocking out *ATOH1* resulted in a strong reduction in tuft cell frequency, while IL-4/IL-13 induced tuft cell proliferation was not affected (Extended Data Fig. 5h-i). Analysis of tuft cell-state specific markers did not point towards the loss of a specific subset (Extended Data Fig. 5j). Thus, *ATOH1* appears to be instrumental in a developmental phase towards tuft cells, but dispensable for tuft cell maintenance. Taken together, we identified four tuft cell states, which developed in a *POU2F3*- and *TCF7*-dependent manner, while IL-4/IL-13-induced tuft cell expansion depended on *HMX2*.

### Single tuft cells give rise to organoids containing all epithelial lineages

Given the tuft cell capacity to expand, we next explored their organoid-forming potential. We purified S/G2/M^+^ tuft cells and non-cycling tuft cells from IL-4 and IL-13 treated FUCCI organoids (Extended Data Fig. 6a). Small spheroids appeared from sorted single KIT^+^ cells at day 3 when cultured in expansion medium (Fig. 4a). Following passaging, tuft cell-derived organoids appeared larger than non-tuft cell-derived organoids (Fig. 4b). Similar results were obtained with sorted organoid-derived single AVIL^+^ cells and adult tissue-derived KIT^+^ cells (Extended Data Fig. 6b-c), albeit at a lower frequency, likely as it contained all tuft cell-states rather than proliferating tuft cells only. Of note, following seeding, AVIL^+^ cells rapidly downregulated tuft cell-specific genes such as *AVIL* and upregulated stem cell markers such as *LGR5* (Extended Data Fig. 6d).

**Fig. 4.**
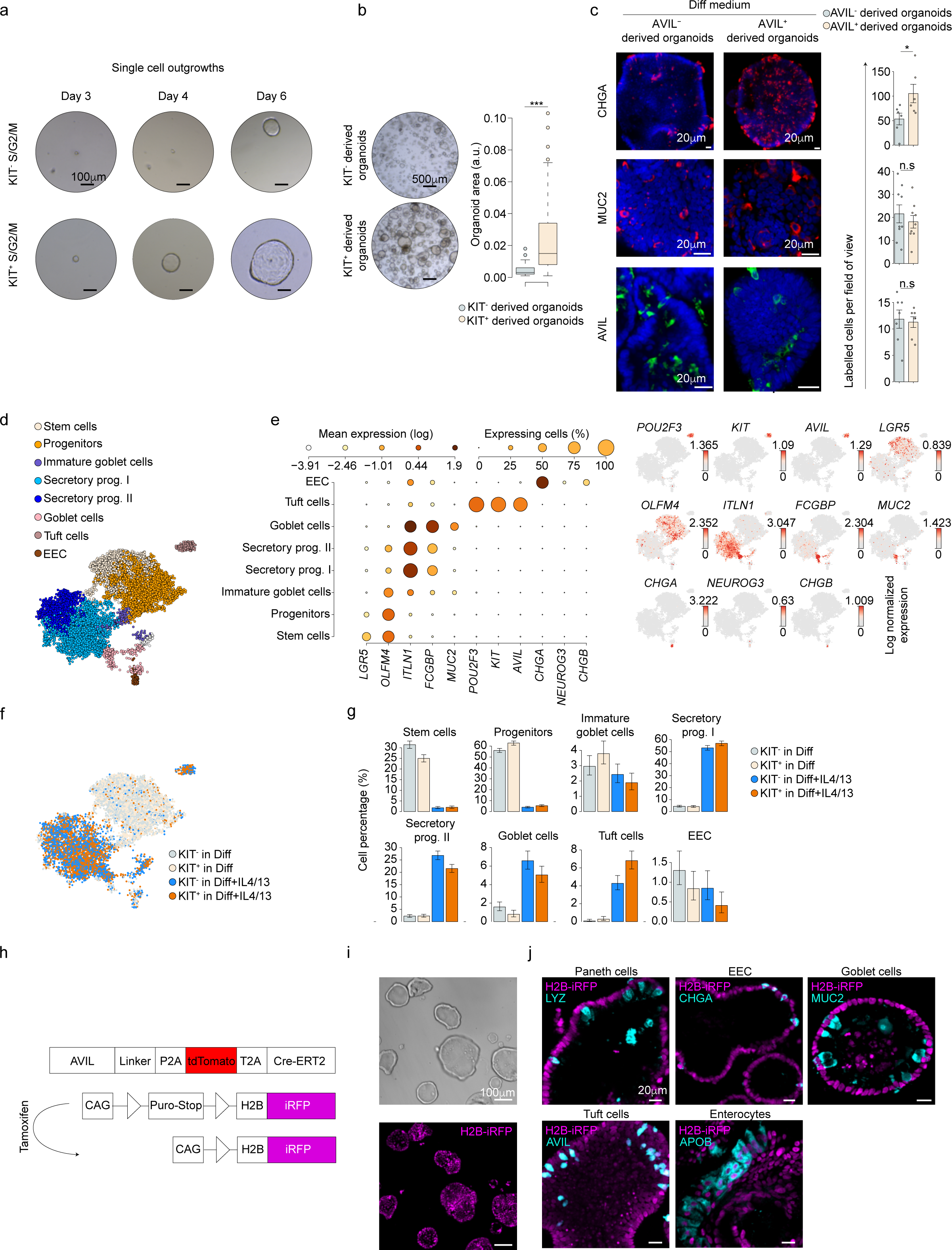
Single tuft cells give rise to organoids that contain all epithelial lineages. **a,** Organoid outgrowth of single S/G2/M phase KIT^+^ and KIT^-^ cells sorted from ileum FUCCI reporter organoids. Sorted single cells were cultured in standard human intestinal expansion medium. **b,** Representative images (left) and area quantification (right) of first passage organoids derived from **(a)**. 47 individual organoids per condition were quantified, shown one of three independent experiments on donor 1 (see Supplementary Information Fig. 4). Box plots present the third quartile (top of the box), median (center lines) and first quartile (bottom of the box) of measurements. The whiskers represent 1.5 times the interquartile range from the top or bottom of the box. Two-tailed Student’s t-test. **c,** Representative images of depicted markers (left) and cell number quantification (right) representing different intestinal epithelial cell types in AVIL^-^ and AVIL^+^ cell-derived organoids at passage 1. Each dot is an individual organoid. Three independent experiments were performed on 2 donors (donor 1 is shown; for donor 2, see Supplementary Information Fig. 4). Error bars indicate SE. Two-tailed Student’s t-test. **d,** Metacell 2D-projection of 10,311 single cells isolated from passage 1 KIT^-^ or KIT^+^ cells-derived organoids, differentiated with or without IL-4/13 supplementation for 7 days. Cells are colored by annotation to cell subsets. **e,** Expression of selected lineage markers across annotated cell types as in **(d)**. Left: Dot color relates to mean expression values and dot size to fraction of expressing cells; right: dot color indicates log normalized expression. **f,** Same as (d), cells are colored by medium condition and by identity of organoid founder cells (KIT^-^ or KIT^+^ cells). **g,** Distribution of the different annotated cell types across medium condition and founder cell. Error bars indicate 95% confidence in binomial estimation of the mean. **h,** Schematics of AVIL lineage tracing approach in human ileum organoids **(i-j)**. **i,** Images of AVIL-lineage tracing organoids, derived from sorted AVIL- tdT^+^iRFP^+^ cells (as in Extended Data Fig. 7d-e, day0); **j,** Representative images of markers of intestinal epithelial lineages in traced organoids differentiated for 5 days in differentiation media. **i-j**, n= 3 independent experiments on donor 1. Diff: human tuft cell differentiation medium; tdT: tdTomato; EEC: Enteroendocrine cells; SE: standard error. * P < 0.05, *** P < 0.001.

In expansion medium and following differentiation, either with or without IL-4 and IL-13, tuft cell (as determined by expression of KIT or AVIL)- and non-tuft cell-derived organoids showed somewhat similar expression levels of genes that define absorptive enterocytes (*FABP1*), goblet cells (*MUC2*), tuft cells (*AVIL*), and stem cells (*LGR5*) (Extended Data Fig. 6e). However, tuft cell-derived organoids expressed higher levels of the entero-endocrine cell (EEC) marker *CHGA* in differentiation medium without IL-4 and IL-13, which was subsequently confirmed by histological analysis (Fig. 4c).

This observed divergence in CHGA expression between tuft cell- and non-tuft cell-derived organoids normalized following further passaging (Extended Data Fig. 6f-g), indicating that the balance between the distinct epithelial subtypes is reestablished over time. In agreement with this, no differences in cell frequencies between passaged tuft cell and non-tuft cell-derived organoids were detected by single-cell RNA-seq analysis (Fig. 4d-g).

This transient effect on *CHGA* expression could not be explained by tuft cells expressing the *CHGA* gene themselves, as WT organoid-derived AVIL^+^ purified cells did not express *CHGA*, while the AVIL^-^ population did (Extended Data Fig. 6h). Indeed, imaging, as well as flow cytometry analysis of differentiated organoids confirmed the mutually exclusive expression of AVIL against either Chromogranin A or Chromogranin B (Extended Data Fig. 6i-k). Interestingly, *POU2F3*-deficient organoids showed reduced *CHGA* expression and protein levels in tuft cell differentiation medium (Extended Data Fig. 6l-m), suggesting the involvement of tuft cells in developing and/or maintaining EEC frequency. Moreover, culture conditions that are known to enhance EEC development and maturation^30^ induced *CHGA* expression only in WT organoids, and not in *POU2F3^-/-^* organoids (Extended Data Fig. 6n).

We formulated two alternative explanations to *CHGA* dependence on tuft cells: either tuft cells act as support cells for EEC development in a paracrine manner, or tuft cells are precursors to EECs. To address this question, we mixed cells from H2B-iRFP-labeled WT organoids (WT^red^) and *POU2F3^-/-^* organoids, forming mosaic organoids (Extended Data Fig. 7a-b). Following differentiation, *CHGA*, *SST*, and *TPH1* expression was enriched in WT^Red+^ cells, but low expression in the *POU2F3^-/-^* cells (Extended Data Fig. 7c), suggesting a paracrine effect. We further implemented a CRISPR Cas9-mediated organoid lineage tracing strategy (Fig. 4h). Here, Cre^ERT2^ is under the control of *AVIL* expression, driving the excision of a stop-sequence that activates a viral promotor-induced H2B-iRFP construct, following tamoxifen administration (Fig. 4h). We observed expression of iRFP^+^ cells that lost AVIL expression, representing tuft cell progeny (Extended Data Fig. 7d-e). We then proceeded to analyze organoids derived from single AVIL^+^iRFP^+^ cells (Fig. 4i, Extended Data Fig. 7d-e). These fully iRFP-labeled organoids contained all major intestinal epithelial lineages, including *CHGA*-expressing EECs (Fig. 4j, Extended Data Fig. 7f). Thus, in agreement with the organoid-forming potential of single tuft cells shown above, AVIL^+^ cells have stem cell-like properties.

### IL-4 signaling enhances a regenerative response driven by tuft cells

We found genes associated with intestinal stem cells to be highly expressed by all tuft cells. These included *ASCL2*, *BMI1*, *SOX4*, but not *OLFM4* and *LGR5*^31^(Fig. 5a, Extended Data Fig. 8a). A few stem cell-associated genes, such as *SMOC2* and *TACSTD2*, were upregulated upon IL-4 and IL-13 exposure, and were indeed upregulated in tuft-3/tuft-4 states (Fig. 5a). Furthermore, organoid-derived tuft cells expressed genes previously associated with fetal gut-like stem cells, facultative stem cells, or revival stem cells, including *TACSTD2*, *MEX3A, PROX1,* and *ANXA1* ^13,16,17^, of which the two latter were also expressed in primary tissue (Fig. 5a, Extended Data Fig. 8a-b), suggesting a role for tuft cells in epithelial renewal following mechanical injury or other insults.

**Fig. 5.**
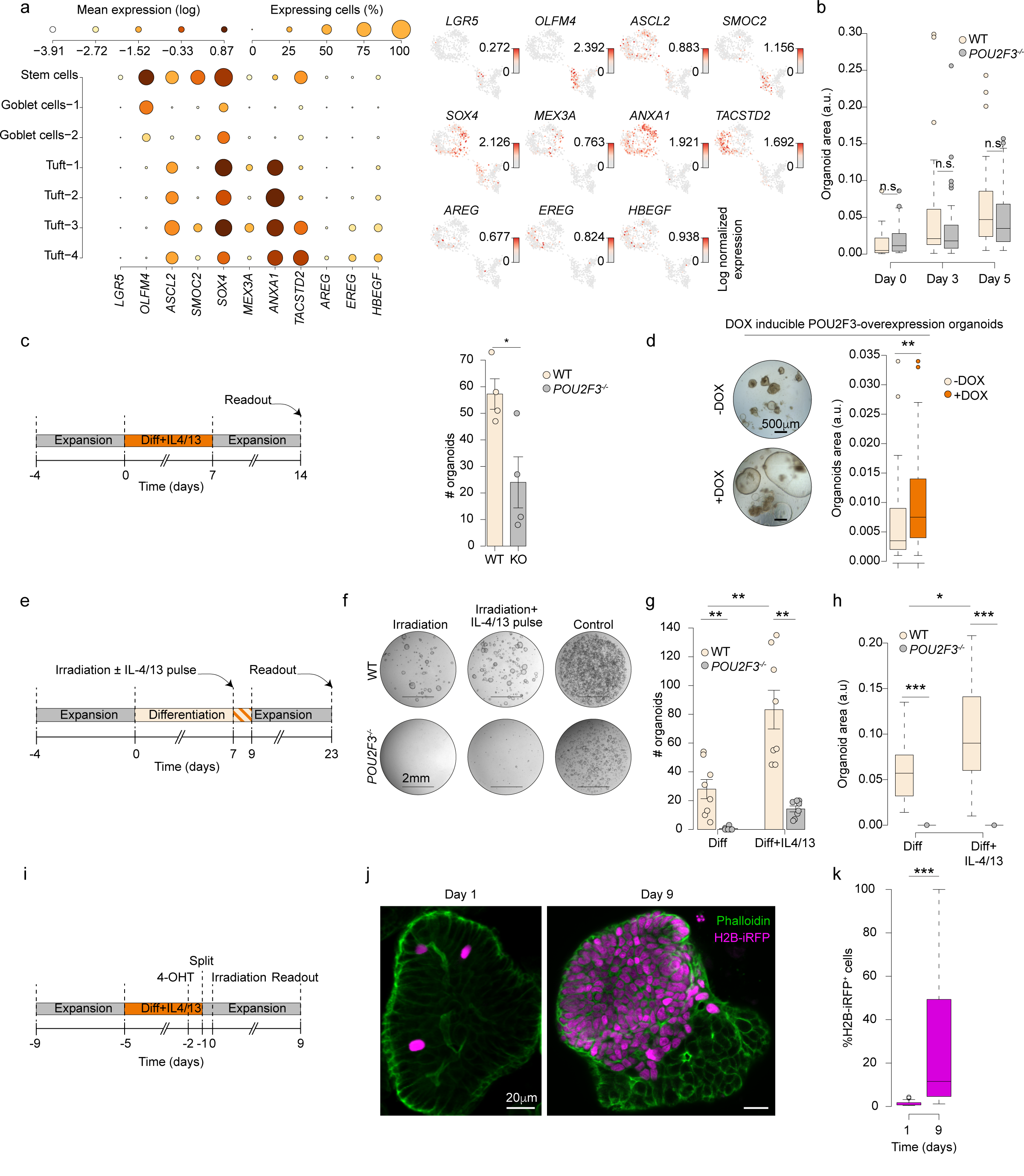
IL-4 signaling enhances the regenerative response in tuft cells. **a,** Gene expression profiles of selected genes associated with adult and repair-induced stem cells from scRNA-seq dataset as in Fig. 3b. Left: Dot color relates to mean expression values and dot size relates to fraction of expressing cells; right: dot color indicates log normalized expression. **b,** Area of organoids cultured in human intestinal expansion medium following passaging at different time points (days). Shown is one of 3 independent experiments on donor 1, 35-51 individual organoids per condition were quantified. Two-tailed Student’s t-test. **c,** Schematics (left) of experimental set-up and quantification (right) of WT and *POU2F3*^-/-^ organoid numbers. Organoids were differentiated for 7 days in tuft cell medium supplemented with IL-4/IL-13, passaged, and cultured for 7 days in human intestinal expansion medium. Each dot is one well. 3 independent experiments were performed on 2 donors (donor 1 is shown; for donor 5, see Supplementary Information Fig. 5). Error bars indicate SE. Two-tailed Mann-Whitney test. **d,** Images (left) and quantification of organoid area (right) from DOX^-^ and DOX^+^ organoids, containing a doxycycline-inducible POU2F3 overexpression construct. Each dot is an individual organoid, n = 46-50 organoids per condition (right). Experiments were performed on 2 donors (donor 2 is shown, for donor 5 see Supplementary Information Fig. 5). Two-tailed Student’s t-test. **e,** Schematics of experimental set-up for **(f-h)**. Following 7 days differentiation in tuft cell medium, organoids were irradiated (5/6 Gy) or not (control), and cultured in tuft cell differentiation medium, with or without a 2 days IL-4/IL-13 pulse. Then, they were passaged and cultured in human intestinal expansion medium for 14 days. **f-h,** Representative images **(f)**, and quantification of organoid numbers **(g)**, and organoid areas **(h)** from organoids as in **(e)**. Experiments were performed on 3 donors (donor 1-2, 4). **f**, donor 2 is shown, for donors 1, 4 see Supplementary Information Fig. 5. **g,** Each dot is one well, pooled from donor 2, 4. Error bars indicate SE. Two-tailed Mann-Whitney test. **h,** 17 individual organoids per condition were quantified. Two-tailed Student’s t-test. **i,** Schematics of experimental set-up for **(j)**–**(k)**. AVIL lineage tracing organoids were differentiated in tuft cell medium with IL-4 IL-13 for 4 days, exposed to 1 μ M 4-OHT for 20 hours, split, and irradiated at 9 Gy. **j**-**k**, Representative images **(j)** and quantification **(k)** of traced cells in organoids at day 1 and day 9 after irradiation. Results are pooled from 3 independent experiments on donor 1, 68 (day 1) or 76 (day 9) organoids were quantified. Two-tailed Student’s t-test. **b, d, h, k,** Box plots present the third quartile (top of the box), median (center lines) and first quartile (bottom of the box) of measurements. The whiskers represent 1.5 times the interquartile range from the top or bottom of the box. Diff: human tuft cell differentiation medium; WT: wildtype; SE: standard error. * P < 0.05, ** P < 0.01, *** P < 0.001.

Mechanical passaging of organoids as a model to induce stress and damage did not show differences in organoid outgrowth dynamics or size between wildtype- and *POU2F3*-deficient organoids (Fig. 5b). However, mechanical disruption of differentiated organoids, followed by expansion, showed a diminished expansion potential in *POU2F3*^-/-^ compared to WT organoids (Fig. 5c). These findings suggested a compromised regenerative capacity in organoids that lack tuft cells. To substantiate these observations, we transfected organoids with a doxycycline-inducible construct that ectopically expresses *POU2F3* under the control of a viral promoter. Indeed, ectopic *POU2F3* expression resulted in a higher frequency of KIT^+^ cells and high expression of tuft cell-related genes following differentiation (Extended Data Fig. 8c-d), and triggered enhanced organoid area as compared to control after mechanical disruption (Fig. 5d). Fast-cycling cells, including stem cells, are vulnerable to irradiation damage. Hence, we irradiated differentiated organoids (5-6 Gy), followed by a 2-day IL-4 and IL-13 pulse (and a non-pulsed control), after which the organoids were allowed to expand (Figure 5e). After an expansion phase of two weeks, *POU2F3^-/-^*organoids gradually lost their potential to passage or expand (Fig. 5f). Wildtype organoids recovered from differentiation and irradiation, and this effect was stronger in organoids that had been exposed to IL-4 and IL-13 (Fig. 5g-h). To further explore whether tuft cells play a major role in reconstitution following injury, we mechanically disrupted and irradiated organoids harboring the AVIL-lineage tracing construct (Fig. 5i, Extended Data Fig. 8e-f). After a recovery period of 9 days, we detected iRFP labeling of entire organoid segments, as well as an overall increase in labeled cell frequency, further confirming the regenerative capacity of tuft cells in these settings (Fig. 5j-k).

Epithelial renewal depends on the presence of growth factors. Our transcriptional analysis showed an IL-4 and IL-13 induced upregulation of *Epiregulin* (*EREG*), and -to a lower extent- *Amphiregulin* (*AREG*) and *heparin binding EGF* (*HB-EGF*) in tuft-3 and tuft-4 cells (Fig. 5a, Extended Data Fig. 8g). The relevance of these growth factors in the context of injury-induced epithelial regeneration was confirmed by employing mechanically disrupted and irradiated organoids engineered to lack *EREG* (Extended Data Fig. 8h-i). Conversely, recombinant EREG supplementation to differentiated *POU2F3^-/-^* organoids partly restored organoid formation (Extended Data Fig. 8j-l). Thus, these experiments implied that tuft cells respond to the wound-repair associated cytokines IL-4 and IL-13 by a rapid proliferative response, providing growth factors, and dedifferentiation of tuft cells to act as reserve stem cells at the affected niche.

The observed stem cell-like transcriptional profile and regenerative properties of adult tuft cells raised the question whether tuft cells in developing intestines contribute to intestinal formation. Upon direct comparison to adult intestinal tissue, fetal intestines showed an increased frequency of KIT^+^ cells (Extended Data Fig. 9a). Fetal KIT^+^ cells exhibited a tuft cell signature on par with adult and pediatric intestinal tuft cells as determined by single cell sequencing, and did not express higher levels of genes associated with a stem cell phenotype (Extended Data Fig. 9b-d). Similarly, histological analysis using an antibody against AVIL confirmed the characteristic tuft cell shape (Extended Data Fig. 9e), and revealed their scattered presence throughout the villus and intervillous zones (Extended Data Fig. 9f). Cell-sorting and subsequent culturing of KIT^+^ and KIT^-^ cells from primary fetal intestinal specimen further revealed that these fetal tuft-like cells displayed potent organoid-forming potential, by comparison to non-tuft cells (Extended Data Fig. 9g-i). The presence of tuft cells in fetal tissues suggest that these cells may be involved in the expansion of the intestinal epithelium during the fetal stages.

## DISCUSSION

Our study of human intestinal tuft cells reveals four distinct tuft cell states, with differential functions. All four states maintain a shared core tuft cell gene expression profile.

Generally, regenerative responses are accompanied by increased IL-4 and IL-13 levels^32^. Our data suggest that previously reported reserve/revival stem cells in mice^17,18,33^ have many features in common with tuft cells described here. Indeed, tuft cells express markers linked to these facultative stem cells, such as *TACSTD2*, *ANXA1*, *BMI1*, and *CLU*. This notion is further supported by a reinterpretation of early studies in mice that linked Dclk1-expressing cells, a then unappreciated murine tuft cell marker, to proliferation in the context of a putative cancer-inducing stem cell population^34,35^. Indeed, in a study employing a BAC-Dclk1::CreERT– dependent strategy to lineage-trace Dclk1-positive cells, it was observed that a small subset of Dclk1^+^ cells could contribute to tissue regeneration upon damage as assessed by lineage tracing *in vivo*. In the same study, it was shown that Dclk1^+^ cells could grow out in short-term 3D culture as organoids, expanding over a 1–2-week period^36^. From the perspective of the current study, we would interpret these combined data to indicate an evolutionary conserved capacity of tuft cells to act as reserve/revival stem cells in the context of intestinal injury.

Taken together, the observations in this study may have implications for regenerative medicine, opening up strategies for therapeutic interventions involving interleukins such as IL-4 and IL-13. Although the current study is focused on the intestinal tract, tuft cells are widely present in other endoderm-derived tissues^37^. It would be of interest to study the potential regenerative roles of tuft cells in other tissues, such as the bile ducts, urinary tract, and airways.

## AUTHOR CONTRIBUTIONS

L.H. designed and performed experiments, and interpreted results. J.B. conceptualized and supervised the project, designed, and performed experiments, interpreted the results, and wrote the manuscript. A.G. interpreted the results, analyzed the data and wrote the manuscript. M.G. assisted with base-editing and knockout strategies in organoid lines. D.K. performed and analyzed live cell imaging, generated and provided several constructs. G.A.B. generated and provided constructs G. S. performed experiments, and analyzed single-cell data. H.B. performed immunohistochemistry experiments. L.L. provided constructs. M.Z. supervised and supported irradiation experiments. H.C. supervised the project, interpreted the results, and wrote the manuscript.

## ACKNOWLEDGEMENTS

We thank Esther Siteur, Kees Weijer, and Stichting Bloemenhove clinic for providing fetal intestinal material, Theodore Grenier for providing adult intestinal material. Talya Dayton for providing paraffin blocks of human intestinal material. L.H. acknowledges financial support from the China Scholarship Council program (201906210081), J.B. acknowledges financial support from ZonMw Veni fellowship (09.150.161.810.107) and Dutch Lung Fund grant (4.2.18.237). A.G acknowledges financial support from the EMBO fellowship (ALTF 112-2022).

## METHODS

### Fetal and adult tissues

Human fetal tissue was obtained from elective abortions at the Stichting Bloemenhove clinic in Heemstede, the Netherlands, upon on the receipt of informed consent. The use of human abortion tissues was approved by the Medical Ethical Committee of the Academic Medical Center, Amsterdam. Gestational age, determined by ultrasonic measurement of the diameter of the skull or femur, ranged from 19 to 21 weeks. Uninflamed ileum, duodenum, and colon were obtained from patients undergoing tumor–resection surgery; ileal, duodenal and colon tissues were collected at an appropriate distance from the tumor. Resection specimen was obtained as residual material after clinical procedures in accordance with the Declaration of Helsinki, the ethical guidelines of the University Medical Centre Utrecht, Utrecht, the Netherlands and Amsterdam UMC, Amsterdam, the Netherlands. This study is compliant with all relevant ethical regulations regarding research involving human participants. Inflammation status was macroscopically deter- mined by the pathologist.

### Generation and Culturing human intestinal organoids

Human intestinal cells were isolated, processed and cultured as previously described^1,2^. Human intestinal organoids were split once a week by mechanic dissociation. In this study, basic culture medium includes advanced Dulbecco’s modified Eagle’s medium/F12 (Gibco) supplemented with 100 U/ml penicillin/streptomycin (Gibco), 10 mM HEPES (Gibco), 1× Glutamax (Gibco), 1× B-27 Supplement (Life Technologies), 1.25 mM N-acetylcysteine (Sigma-Aldrich) and 1% (v/v) recombinant Noggin (U-Protein Express).

Organoids were expanded in human expansion medium as described before^1^. For differentiation toward tuft cells, organoids were washed 30 minutes in DMEM+++ at day 4 and the medium is replaced for tuft cell differentiation medium (diff): 0.5 nM Wnt surrogate (U-Protein Express), 20% (v/v) R-spondin1 (condition medium), 50 ng/ml recombinant human EGF (Peprotech), 10 μM Notch inhibitor DAPT (Sigma-Aldrich) in basic culture medium. For tuft cell differentiation with IL-4 and IL-13, EGF was withdrawn, 5 ng/ml human IL-4 (Peprotech) and 5 ng/ml human IL-13 (Peprotech) were supplemented to tuft cell differentiation medium (diff). BMP activation was achieved by withdrawing Noggin and addition of 50 ng/ml BMP-2 (Peprotech) and 50 ng/ml BMP-4 (Peprotech). Mature enterocytes and Paneth cell differentiation in Figure 4j was achieved by using EGF, R-spondin1, BMP-2/4) (ERB medium)^3^ or adding 10 ng/ml human IL-22 (Peprotech) in WENRA^4^ (Wnt/ R-spondin1, EGF, Noggin, ALK4,5,7 inhibitor (A83-01, Tocris) medium respectively. For specific experiments, 10 ng/ml human IL-27 (Peprotech), 10 ng/ml human IL-25 (Peprotech), 10 ng/ml human SCF (Peprotech), 50 ng/ml recombinant human Epiregulin (Peprotech), were used.

For organoids outgrowth experiment in Figure 4a-c, 100 single cells per 10 μl BME were seeded, and replated at day 3, keeping similar seeding density between tuft cell and non-tuft cell conditions.

For irradiation experiments, culture plates were sealed air-tight and irradiated with a single fraction of 5-6 Gy using a linear accelerator (Elekta Precise Linear Accelerator 11F49, Elekta). The plates were positioned on top of 2-cm polystyrene and submerged in a 37°C water bath and radiated from below with the plate being positioned at exactly 100cm from the radiation source. After radiation, medium was changed (tuft cell differentiation medium without EGF, with or without IL-4/IL-13).

For irradiation of AVIL-lineage tracing organoids in Figure 5i, organoids were differentiated 4 days in tuft cell differentiation medium with IL-4 and IL-13, exposed to 1 μM Tamoxifen for 20 h, then the organoids were split, and irradiated at 9 Gy one day after splitting. After splitting and irradiation, the organoids were grown in human intestinal expansion medium.

### Generation of stable genetically modified organoids

Reporters: For generation reporter organoids using a CRISPR-HOT approach as described in^5^, human duodenum, ileum and colon organoids were dissociated into small clumps, washed twice with Opti-MEM (Thermo Fisher scientific) and resuspended in BTXpress solution (BTX). Clumps were resuspended with a targeting plasmid containing a fluorescent protein (Clover, mNeon or tdTomato) which can be linearized at a defined base position by a specific sgRNA and Cas9 (provided from a second plasmid, frame selector plasmid which also encodes mCherry)^6^. These two plasmids were co-electroporated with a plasmid encoding the sgRNA for the *gene* locus (Supplementary Table 2, Addgene #47108, #66939, #66940, #138569, #174092). Following blasticidin selection or cell sorting based on mCherry signal, clones were picked and successful incorporation was confirmed by Sanger sequencing (Macrogen).

Knockouts: For generation knock out organoid lines using base-editing or conventional CRISPR-Cas9, spacer sequences for sgRNAs were cloned as previously described in the empty sgRNA backbone that was a kind gift from Keith Joung (BPK1520, Addgene #65777)^7^. In short, plasmids were amplified using -inverse PCR (Q5, NEB), using primers with overhangs containing spacer sequences. PCR amplicons were subsequently ligated using T4 ligase (NEB) in a reaction with DpnI (NEB) to remove PCR template material. Ligations were transformed into MACH1T1 cells (Thermofisher) and sgRNA identity was confirmed by Sanger sequencing. For electroporation, 2.5 μg sgRNA plasmid (BPK1520, Addgene #65777), 7.5 μg pCMV_AncBE4max_P2A_GFP plasmid (for base-editing, Addgene #112100) or pCAS9- mCherry-Frame +1 plasmid (for conventional CRISPR-Cas9, Addgene #66940), together with 10 μg PiggyBac transposon system (5 μg transposase + 5 μg hygromycin resistance containing transposon)^8^ were co-electroporated into human duodenum, ileum and colon *AVIL*-Clover reporter organoids. *ATOH1*^-/-^ line was generated in human ileum wildtype organoids. After hygromycin selection, subclones were genotyped. Knockout clones were further expanded for the following experiments. The list of gRNAs and primers to genotype can be found in Supplementary Table 2.

FUCCI and overexpression constructs: For generation FUCCI reporter (in wildtype human ileum organoids), *ATOH1* overexpression and H2B-iRFP in human ileum *AVIL*-Clover reporter organoids using transposon system, the FUCCI construct, *ATOH1*-P2A-iRFP670 overexpression plasmid and H2B-iRFP670 construct were cloned into a p2T-based vector^9^ respectively using Gibson Assembly (NEBuilder HiFi DNA Assembly). The FUCCI sequence comprising mCherry-Cdt1-T2A-Geminin-hmAzami-Green was PCR-amplified from a construct provided as a kind gift by Geert Kops (Hubrecht institute, Utrecht). The ATOH1-P2A- iRFP670 overexpression plasmid was cloned using a three-insert Gibson reaction. ATOH1 cDNA (IDT) was first cloned into a backbone vector with GSlinker-P2A-iRFP670, and the complete fragment of ATOH1-GSlinker-P2A-iRFP670-SV40polyA was amplified. The DNA fragment of tight TRE promoter with ATG start codon and 3XFLAG tag was amplified from pCW-Cas9 (Addgene #50661). The DNA fragment of hPGK-PuroR-rTetR was amplified from pCW-Cas9. All three DNA fragments were then cloned in the digested backbone. For continuous expression H2B-iRFP670, the H2B-iRFP670 and IRES-Puromycin sequences were PCR-amplified respectively, then were inserted into the p2T vector cut with NheI (Promega) and SmaI (Promega). Organoid lines were generated by co-electroporation of 5 µg of the respective FUCCI, *ATOH1*-P2A-iRFP670 or H2B- iRFP670 expression construct together with 5 µg mT2TP transposase mediating the tol2-dependent random integration of the expression constructs into the cell genome.

For generation *POU2F3* overexpression in human duodenum, colon organoids using lentivirus system, a gBlock for *POU2F3* was ordered from IDT, which contained a 5’UTR sequence, including an EcoRI site and a Kozak sequence, and 3’ tag sequence instead of the STOP codon, which includes a Gly linker, HA tag, P2A sequence and another EcoRI site. This gBlock was cloned into a pJet vector and an EcoRI restriction enzyme cloning step was done to introduce the *POU2F3* sequence into pLX vector, which enable the doxycycline-induced *POU2F3*-P2A-tdTomato expression^10^. Organoids were lentivirally transduced as described before^11^.

Lineage tracing: Human ileum organoids were targeted with a tdTomato and CreERT2 sequence fused with the gene encoding for Advillin using a CRISPR-HOT approach as described in^5^ (tdTomato-T2A-CreERT2 to the AVIL locus separated by a P2A sequence to obtain AVIL-P2A-tdTomato-T2A-CreERT2). In parallel, using mT2TP transposase-based random integration, a CAG promoter-driven loxp-flanked puromycin resistance containing a stop codon with a downstream H2B-iRFP670 was introduced (CAG-loxp-puromycin-stop-loxp- H2B-iRFP670).

### Transmission electron microscopy

Organoids were chemically fixed for 3 hours at room temperature with 1.5% glutaraldehyde in 0.067 M cacodylate buffered to pH 7.4 and 1 % sucrose. Samples were washed once with 0.1 M cacodylate (pH 7.4), 1 % sucrose and 3x with 0.1 M cacodylate (pH 7.4), followed by incubation in 1% osmium tetroxide and 1.5% K4Fe(CN)6 in 0.1 M sodium cacodylate (pH 7.4) for 1 hour at 4 ᵒC. After rinsing with MQ, organoids were dehydrated at RT in a graded ethanol series (70, 90, up to 100%) and embedded in Epon polymerized for 48h at 60 ᵒC. Ultrathin sections of 60 nm were cut using a diamond knife (Diatome) on a Leica UC7 ultramicrotome, and transferred onto 50 Mesh copper grids covered with a Formvar and carbon film. Sections were post-stained with uranyl acetate and lead citrate.

All TEM data were collected autonomously as virtual nanoscopy slide on Tecnai T12 microscopes (Thermo Fisher Scientific) at 120kV using an Eagle camera. Data were stitched, uploaded, shared and annotated using Omero and PathViewer. The final pictures were directly acquired at the microscope in a manual standard way. The final pictures were directly acquired at the microscope in a manual standard way using the Eagle camera at 4kx4k.

### RNA isolation and quantitative PCR

Organoid RNA was isolated using RNAeasy kit (QIAGEN), following the manufacturer’s protocol. Quantitative PCR (qPCR) analysis was performed biological and technical replicates as described before^12^. Primers were listed in Supplementary Table 3.

### Flow cytometry

Organoids were dissociated into single cells using TrypLE (TrypLE Express, Life Technologies) with 10 μM Rho kinase inhibitor (Abmole) in 37°C and mechanical disruption by pipetting every 5 minutes. Cells were stained 30 mins with antibody, then were analyzed using a BD LSR Fortessa X20 4 laser (BD Biosciences) based on fluorescence levels. For single cell RNA sequencing, single cells were sorted on FACSFusion (BD Biosciences) and collected in 384- well plates with ERCC spike-ins (Agilent), reverse transcription primers and dNTPs (both Promega). Single cell sequencing was performed according to the Sort-seq method^13^.

For organoids staining, PE anti-human CD117 antibody (KIT; Biolegend, 313204), or Biotin anti-human CD117 (Biolegend, 313208) and Brilliant Violet 421™ Streptavidin (Biolegend, 405226) were used in some experiments. For sort from fresh human fetal/colon intestine tissue, Alexa Fluor 488 anti-human CD326 (Epcam; Biolegend, 324210), APC/Cy7 anti-human CD45 (Biolegend, 304014), PE anti-human CD117 antibody (KIT; Biolegend, 313204) were used. For cell multiplexing oligo labeling, organoids were digested into single cells using TryplE (Thermo Fisher), washed three times with ice-cold PBS + 10% FBS and incubated 15 mins at RT with 100 μl Cell Multiplexing Oligo. After wash, 8000 live cells (DAPI^-^ cell) per condition were sorted into collection tube, were subjected to droplet-based scRNA-seq using the 10x Genomics platform.

### Histology and immunostainings

For immunostainings, sections of formalin-fixed, paraffin embedded human colon and ileum tissue were obtained from resections performed at the University Medical Center Utrecht, the Netherlands. Anonymized archival pathology material was used according to the guidelines of the UMC Utrecht’s Research Ethics Committee^14^. The human intestine tissues were fixed 2 h at room temperature in 4% formalin, embedded in paraffin and stained as described previously^15^. Rabbit anti-Advillin (1:500, Sigma-Aldrich, HPA058864), mouse anti-KI67 (1:4000, monosan, MONX10283) followed by goat-anti-rabbit or goat-anti-mouse conjugated to horseradish peroxidase (1:2000, Dako, P0448) and then visualized (VS200 slide scanner, Olympus-lifescience).

Whole mount staining of organoids was performed as described before^16^. In brief, organoids were removed from the BME, then were fixed 2 h at room temperature in formalin. Next, the organoids were permeabilized using 0.1% Tween 20 (Sigma-Aldrich) in PBS for at least 15 min and blocked at least 1 h in 0.1% Triton X-100 (Sigma-Aldrich), 1 g/l BSA (Sigma-Aldrich) in PBS. Primary antibodies used were rabbit anti-AVIL (1:600, Sigma-Aldrich, HPA058864), goat anti-GFP (1:600, Abcam, ab6673), mouse anti-KI67(1:600, Abcam, ab16667), rabbit anti-chromogranin A (1:600, labned.com, LN1401487)), rabbit anti-mucin 2 (1:200, santa cruz biotechnology, sc-15334), rabbit anti-APOB (1:100, Novus biologicals, NBP2-38608), rabbit anti-Lysozyme antibody (1:100, GeneTex, GTX72913), rabbit anti-Vimentin (1:100, Cell Signaling technology, 5741S), mouse anti-TM4SF4 antibody (1:100, Sigma-Aldrich, sc- 293348), APC Mouse anti-human CD274 (BD Pharmingen, 563741). Organoids were incubated with Phalloidin–Atto 647N (Sigma-Aldrich, 65906) or the corresponding secondary antibodies Alexa488-, 568- and 647-conjugated anti-goat, anti-rabbit and anti-mouse (1:1000, Molecular Probes) in blocking buffer containing 4ʹ,6-diamidino-2-phenylindole (DAPI, 1:1000, Invitrogen). Sections were embedded in fructose–glycerol clearing solution Image analysis was performed using ImageJ and Imaris software.

Some images were obtained using a Zeiss LSM880 confocal microscope with Airyscan (Carl Zeiss) and a LCI Plan-Neofluar 63x/N.A. 1.3 water immersion objective (Carl Zeiss) at a voxel resolution of 0.04 µm (x/y) to 0.19 µm (z). Images were deconvoluted using the Zen Black (Carl Zeiss)-inbuild Airyscan post-processing module. Images were processed (Gauss filtering) using Fiji and rendered in Imaris (Andor Technology).

STED super resolution microscopy was performed using a Leica STELLARIS 8 STED microscope using a HC PL APO CS2 100x/N.A. 1.40 oil objective. Organoids expressing AVIL- Clover were fixed, stained with DAPI and phalloidin-atto647N and mounted on 0.16–0.19 mm thick cover glasses (Glaswarenfabrik Karl Hecht GmbH & Co KG) in ProLong Gold Antifade Mountant (Thermo Fisher Scientific). The tuft cell reporter and DAPI signal was recorded with confocal resolution and tuft cells identified. The actin cytoskeleton was visualized with super resolution microscopy with a pixel resolution of 10 x 10 nm. To do so, phalloidin-atto647N was illuminated with 647 nm excitation and 775 depletion lasers. The signal was recorded with a line averaging of 16, a dwell time of 0.69 µs.

For clustering analysis, the surface of organoids in 3D image stacks was projected into 2D using the *LocalZProjector* Fiji plugin based on DAPI-stained nuclei and tuft cells were segmented using the ITK-SNAP software. For the clustering analysis, a custom-made Matlab script was used. Inter-tuft cell distance was measured based on the Euclidean distance of their centroid position and expressed as a multiple of the average cell distance. A density map was computed, and a contour plot was generated using the *imcontour* function. Clusters were identified within an 8-cell-distance and analyzed using the Matlab package “Distance-based clustering of a set of XY coordinates” (Yann Marcon (2023), made available through Matlab Central File Exchange).

### Live-cell imaging of human intestinal organoids

Live imaging experiments were performed on a Leica SP8 confocal laser scanning microscope equipped with Argon laser and White Light Laser at 37°C and 5% CO2 using a Leica 20x/NA0.7 Air objective. Images were acquired in a line-sequential mode separating the fluorophore recordings with minimal spectral overlap with a final pixel resolution of 1.65 pixels/µm, an axial resolution of 1.4 µm and a time interval of 9 min.

Long-term live imaging was performed using a LS1 Live light sheet microscope (Viventis Microscopy) using a Nikon 25X NA 1.1 water immersion objective at a magnification of 18x. Organoids were mounted on a single-chamber sample holder one day before the start of imaging. A position-specific alignment of the light sheets with a thickness of 2.2 µm was done. The samples were imaged with a 488 nm and 561 nm illumination to visualize Clover- and tdTomato-based reporters with a time interval of 10 min at 37°C and 5% CO2.

The Post-acquisitional analysis was done with custom-made Fiji-script.

### Single cell RNA sequencing analysis

Single cell RNA-seq libraries of organoid derived material, and KIT^+^ enriched cells from fetal and adult intestines were sequenced on an Illumina NextSeq platform, at a median sequencing depth of 49,861 reads per cell. Reads were mapped to a human genome (hg38) integrated with the Clover transcript using STAR (version 2.7.8a), reads with multiple mapping positions were excluded. Reads were associated with genes if they were mapped to an exon. Reads were demultiplexed and collapsed into UMI tables using umi_tools (version 1.1.1) allowing up to one hamming distance of the cell barcode. Cells with less than 500 UMI, or with more than 40% mitochondrial genes were excluded from analysis.

All analysis was performed in R. We used the MetaCell package^17^ to analyze all scRNA-seq data collected in this study. Default parameters were used unless otherwise stated. We derived a metacell cover of DAPI^-^/AVIL-Clover^+^ and DAPI^-^ epithelial cells from human ileal organoids. Mitochondrial genes and the highly variable immunoglobulin genes (IGH, IGK and IGL prefixes) were removed from the UMI tables. Gene features for metacell covers were selected using the parameter Tvm=0.1, total umi > 10, and more than 3 UMI in at least 3 cells. We filtered the list of gene features used for metacell analysis from genes associated with cell cycle, immediate stress response and gene modules inducing strong patient-specific biases. To this end, we first identified all genes with a correlation coefficient of at least 0.13 for one of the anchor genes TOP2A, NKI67, PCNA, MCM4, UBE2C, STMN1, FOS, EGR1, IER3, FOSB,

HSPA1B, HSPA1A, HSP90AA1, and DNAJB1. We then hierarchically clustered the correlation matrix between these genes (filtering genes with low coverage and computing correlation using a down-sampled UMI matrix) and selected the gene clusters that contained the above anchor genes. We thus retained 94 genes as features. We used metacell to build a kNN graph, perform boot-strapped co-clustering (500 iterations; resampling 70% of the cells in each iteration), and derive a cover of the co-clustering kNN graph (K=30). Outlier cells featuring gene expression higher than 4-fold than the geometric mean in the metacells in at least one gene were discarded. Detailed annotation of the different tuft and epithelial cell subsets was performed using hierarchical clustering of the metacell confusion matrix. ClusterProfiler^18^ (version 3.14.0) and ChIPpeakAnno (version 3.20.0) were applied to perform gene functional annotation of differentially expressed genes.

scRNA-seq of passage 1 KIT^-^ and KIT^+^ derived organoids was performed using the Chromium Next GEM Single Cell 3’ v 3.1 platform, and sequenced on an Illumina NovaSeq6000 platform. Reads were mapped to the human genome (hg38) and demultiplexed using cellranger (version 7.1.0). Recovered cellplex barcodes were used to assign single cells to experimental batches. Single cells with less than 64 UMI of a specific cellplex barcodes were discarded from down-stream analysis. Single cells with less than 8-fold UMI count ratio between highest and second highest cellplex barcodes were marked as doublets and discarded from down- stream analysis. Single cells with less than 1,000 genomic UMIs or more than 20% mitochondrial content failed to pass QC and were discarded from further analysis, resulting in 10,311 QC-positive cells.

Clustering of passage 1 KIT^-^ and KIT^+^ derived organoids was performed as stated above. Gene features for the metacell covers were selected using the parameter Tvm=0.1, total umi > 15, and more than 3 UMI in at least 3 cells, resulting in 228 features.

We reanalyzed scRNA-seq data from human primary intestinal tissue^19^. We selected 15,184 single cells from healthy adult small intestine, with >1,000 and <20,000 total UMI for further analysis. Cells were analyzed with the metacell package as previously described to derive a 2D representation of the data for Extended Data Figures 2i and 5a. Otherwise, we used predefined annotations to epithelial cell types. In Fig. 2h and 3m we sampled 500 cells from each cell types out of the total 77,364 healthy adult single cells in that database.

### Quantification and statistics

All experiments were performed in multiple distinct replicates, as indicated in the text and figure legends. All statistical tests were two-tailed, except in fig 1f, 2c and supp fig. 1b, where different growth conditions were assessed for increased tuft cell numbers. We used Student’s t-test for continuous data and Mann-Whitney test for discrete data, and used FDR adjustment to correct for multiple hypotheses.

### Data availability

Organoid and primary tissue single-cell RNA-seq that support the findings of this study, were deposited in the Gene Expression Omnibus (GEO) under accession code GSE233451, and will be made publicly available upon publication.

### Code availability

All the custom code and scripts used to analyze the data and produce the figures will be made available in the github repository of A.G.

---

**Extended Data Fig. 1.**
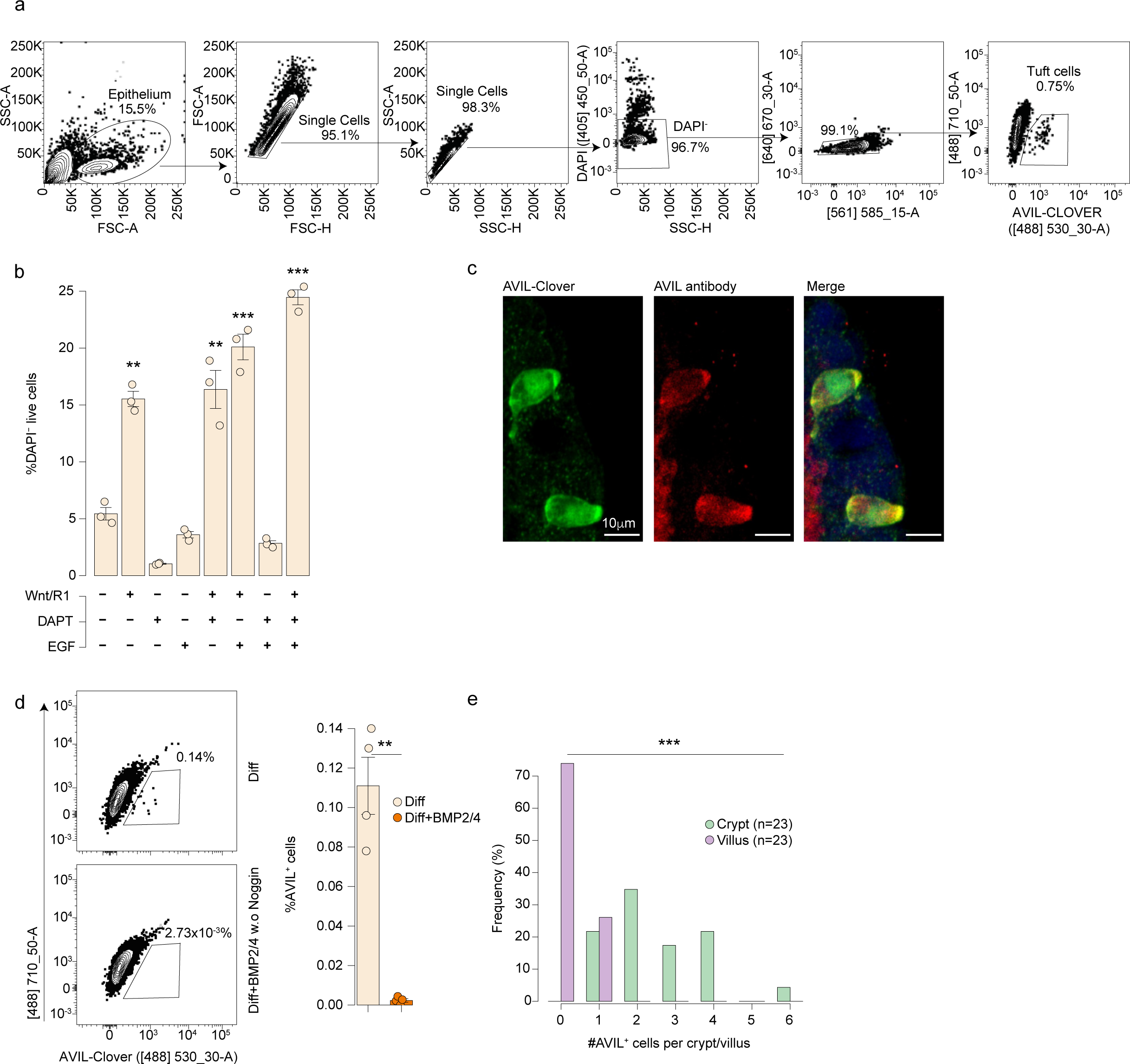
Flow cytometry gating strategies and quantifications of AVIL^+^ cells. **A,** Gating strategy for flow cytometric analysis of AVIL-Clover reporter organoids. **b,** Quantification of DAPI^-^ live cells in ileum organoids differentiated in different regimens. Each dot is a well. One of 3 independent experiments on donor 1 is shown (Supplementary Information Fig. 1). Error bars indicate SE. FDR-adjusted one-sided Student’s t-test against base medium. **c,** Fluorescence images of AVIL-Clover reporter activity (left), AVIL antibody staining (middle), and merged (right). n= 4 donors (donor 1 is shown). **d,** Flow cytometric analysis (left) and quantification (right) of the percentage of AVIL^+^ cells in human ileum AVIL- Clover reporter organoids differentiated for 4 days in tuft cell medium without Noggin, with or without addition of BMP2/BMP4. Each dot is one well. 3 independent experiments were performed on 2 donors (donor 1 is shown; for donor 3, see Supplementary Information Fig. 1). Error bars indicate SE. Two-sided Student’s t-test. **e,** Histogram of AVIL^+^ cell numbers on the crypt/villus axis in human ileum tissue. n = 23 crypt-villus axes from one donor; for 3 additional donors, see Supplementary Information Fig. 1. Error bars indicate SE. χ^2^ test. Diff: human tuft cell differentiation medium; SE: standard error. **P< 0.01, ***P< 0.001.

**Extended Data Fig. 2.**
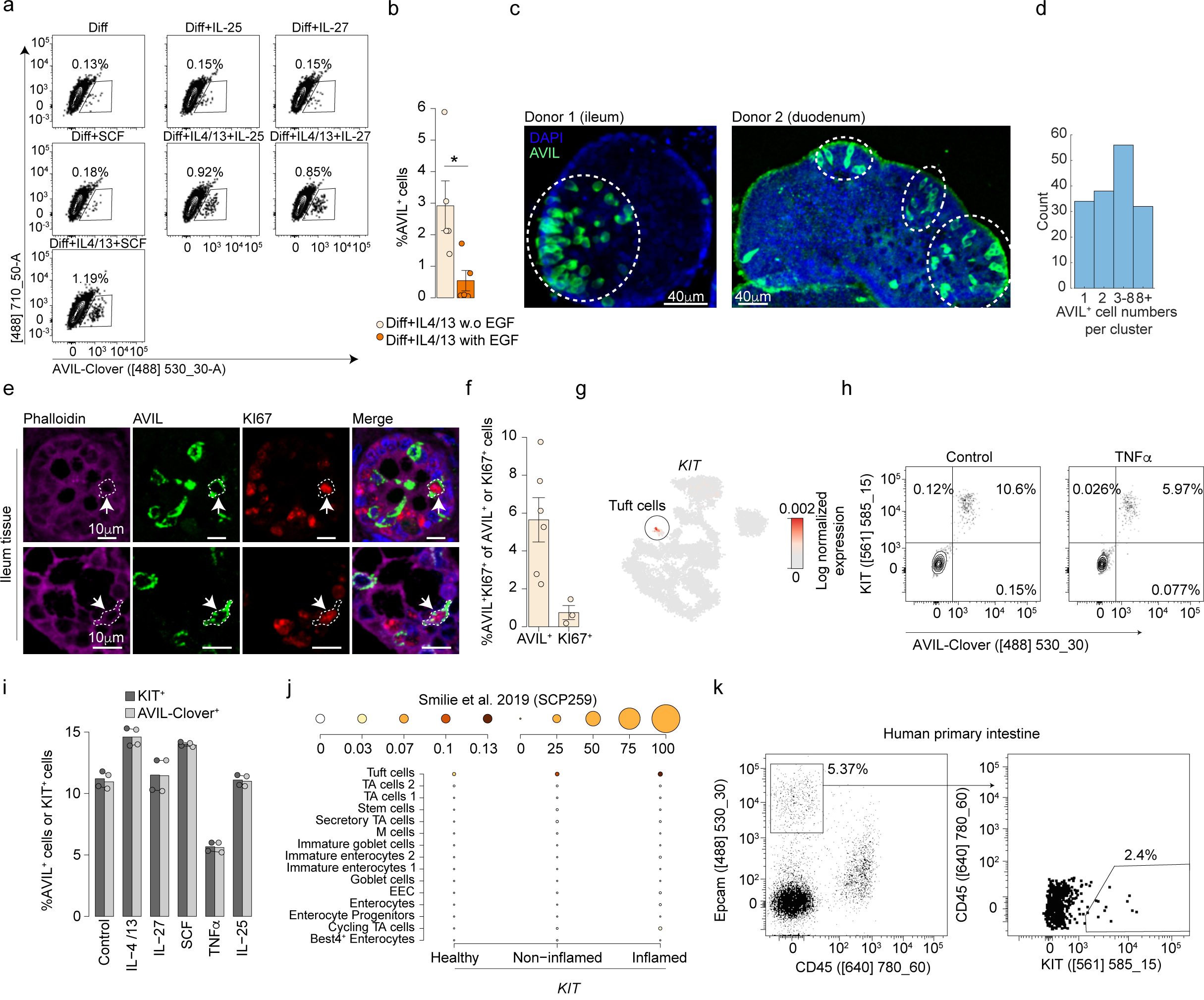
Tuft cell expansion in different conditions. **a,** Representative flow cytometric analysis of AVIL-Clover reporter organoids differentiated for 4 days in tuft cell medium supplemented with indicated recombinant proteins. **b,** Quantification of AVIL^+^ cell frequency in human small intestinal AVIL-Clover organoids differentiated for 4 days in tuft cell medium with IL-4 and IL-13, with or without EGF. Each dot is one well, results are pooled from 3 donors (donor 1-3). Error bars indicate SE. Two-tailed Student’s t-test. **c-d,** Representative images **(c)** and quantification **(d)** of AVIL^+^ cell clusters in IL-4 and IL-13 treated human intestine organoids. **c,** Dashed ellipses indicate clusters of AVIL^+^ cells, (Supplementary Information Fig. 2). **d,** Using an 8-cell distance-based clustering model, each AVIL^+^ cell was assigned to a cluster or remains isolated, and the number of AVIL^+^ cells per cluster were quantified; n= 988 AVIL^+^ cells (30 organoids) pooled from 2 donors (donor 1-2). **e**, Fluorescence images of histological sections of human ileum tissues co-stained for Phalloidin (F-actin, magenta), AVIL (green), KI67 (red) and DAPI (blue). Arrows and dashed lines indicate cells with AVIL and KI67 overlap. n=2 donors. **f**, Quantification of AVIL^+^KI67^+^ cells in histological sections of human ileum tissue. Values indicate AVIL^+^KI67^+^ cell percentage out of the AVIL^+^ or KI67^+^ cells. Each dot is a pool of at least 10 crypts, 1,908 cells in 323 crypts from 3 donors were quantified. Error bars indicate SE. **g**, Log-normalized expression of *KIT*, projected on a Metacell 2D representation of scRNA-seq data of primary human adult small intestine^18^. **h-i**, Representative flow cytometry analysis **(h)** and quantification **(i)** of the KIT^+^ and AVIL^+^ cells in KIT labeled AVIL-Clover organoids triggered with the depicted cytokines for 48 hours, followed by 6 days culturing in tuft cell differentiation medium with IL-4 and IL-13. Each dot is a well. Experiments were performed on 2 donors (donor 1 is shown; for donor 4, see Supplementary Information Fig. 2). **j**, *KIT* expression across epithelial cell types in healthy, IBD non-inflamed, and IBD inflamed human colon tissue^23^. Dot color relates to mean expression values and dot size to fraction of expressing cells. n= 4,428 cells. **k,** Gating strategy of sorting KIT^+^ cells from human adult ileum and colon tissue. n= 2 donors (colon is shown). Diff: human tuft cell differentiation medium; TA: Transit-Amplifying Cells; EEC: Enteroendocrine cells; IBD: Inflammatory Bowel Disease; SE: standard error. *P< 0.05.

**Extended Data Fig. 3.**
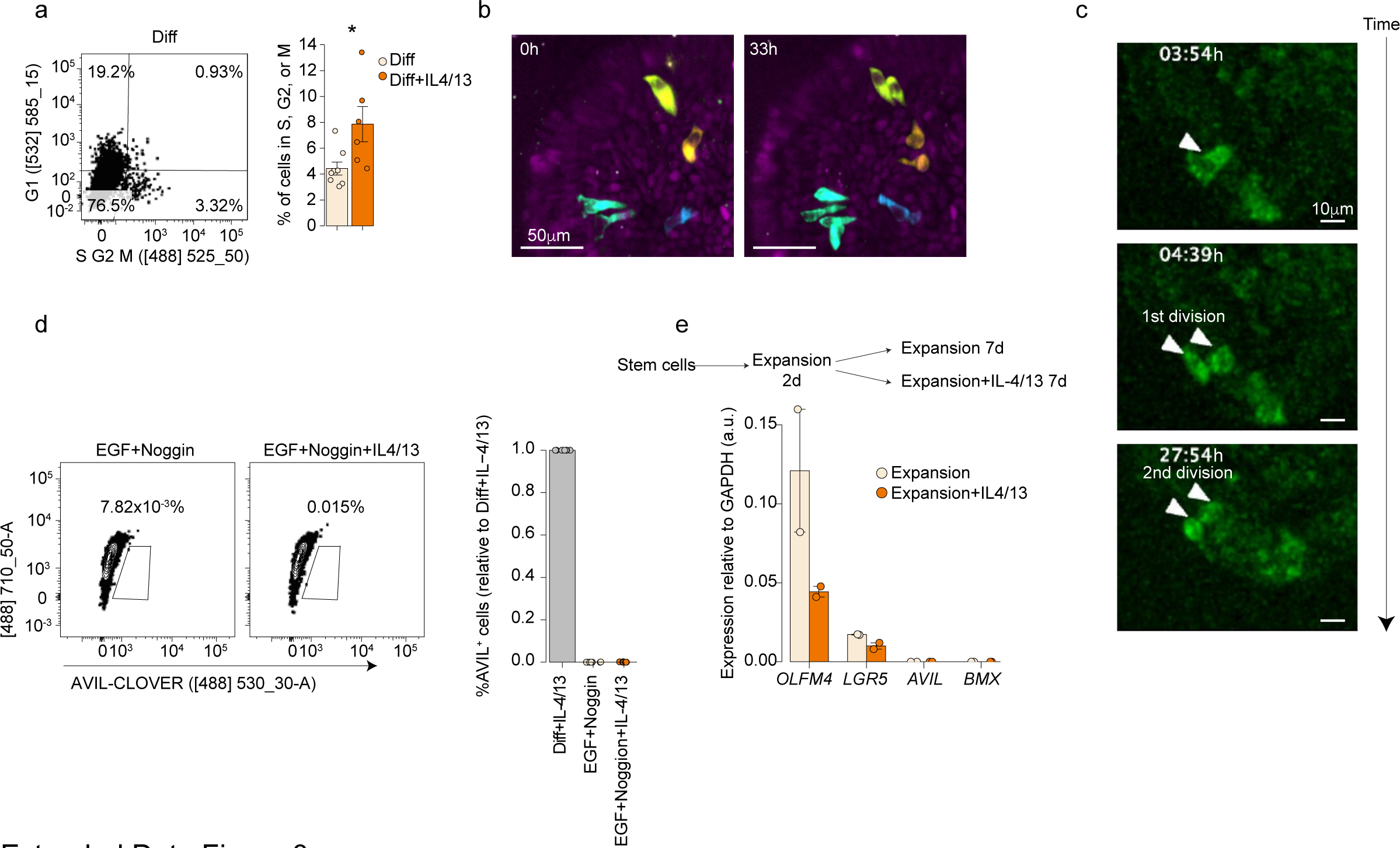
Proliferation dynamics of tuft cells. **a,** Representative flow cytometric analysis (left) and quantification (right) of the S/G2/M phase in KIT^+^ cells in FUCCI reporter organoids differentiated for 3 days in tuft cell medium with or without IL-4/13 treatment. Each dot is one well. Results are pooled from 6 (Diff) or 4 (Diff+IL-4/13) independent experiments. Error bars indicate SE. Two-tailed Student’s t-test. **b-c,** Snapshots of dividing AVIL^+^ cells by live-cell imaging using AVIL-Clover reporter organoids differentiated in tuft cell medium with IL-4/IL-13 for 3 to 5 days **(b)**, or 5 to 8 days **(c)**. **b,** Each dividing AVIL^+^ cell and its progeny are colored differently; **c,** Shown is one tuft cell dividing twice within 24h. **b,** donor 1; **c,** donor 2 (Supplementary Video 3, 5). **d,** Representative flow cytometric analysis (left) and quantification (right) of AVIL^+^ cell frequency in organoids differentiated for 2 days in EGF Noggin medium (tuft cell medium without Wnt, Rospond1 and DAPT), followed by 4 days incubation with or without IL-4/IL-13. Right: each dot is a well, results are pooled from 3 independent experiments from two lines of donor 1. Error bars indicate SE. **e,** qPCR quantification of tuft cell and stem cell genes in single cells sorted from organoids cultured in human expansion medium. Single cells were seeded in expansion medium for 2 days, then cultured for 7 days with or without IL-4/IL-13 treatment. Each dot is a technical duplicate. Experiment was performed on 2 donors (donor 1 is shown; for donor 4, see Supplementary Information Fig. 2). Error bars indicate SE. Diff: human tuft cell differentiation medium; SE: standard error. *P< 0.05.

**Extended Data Fig. 4.**
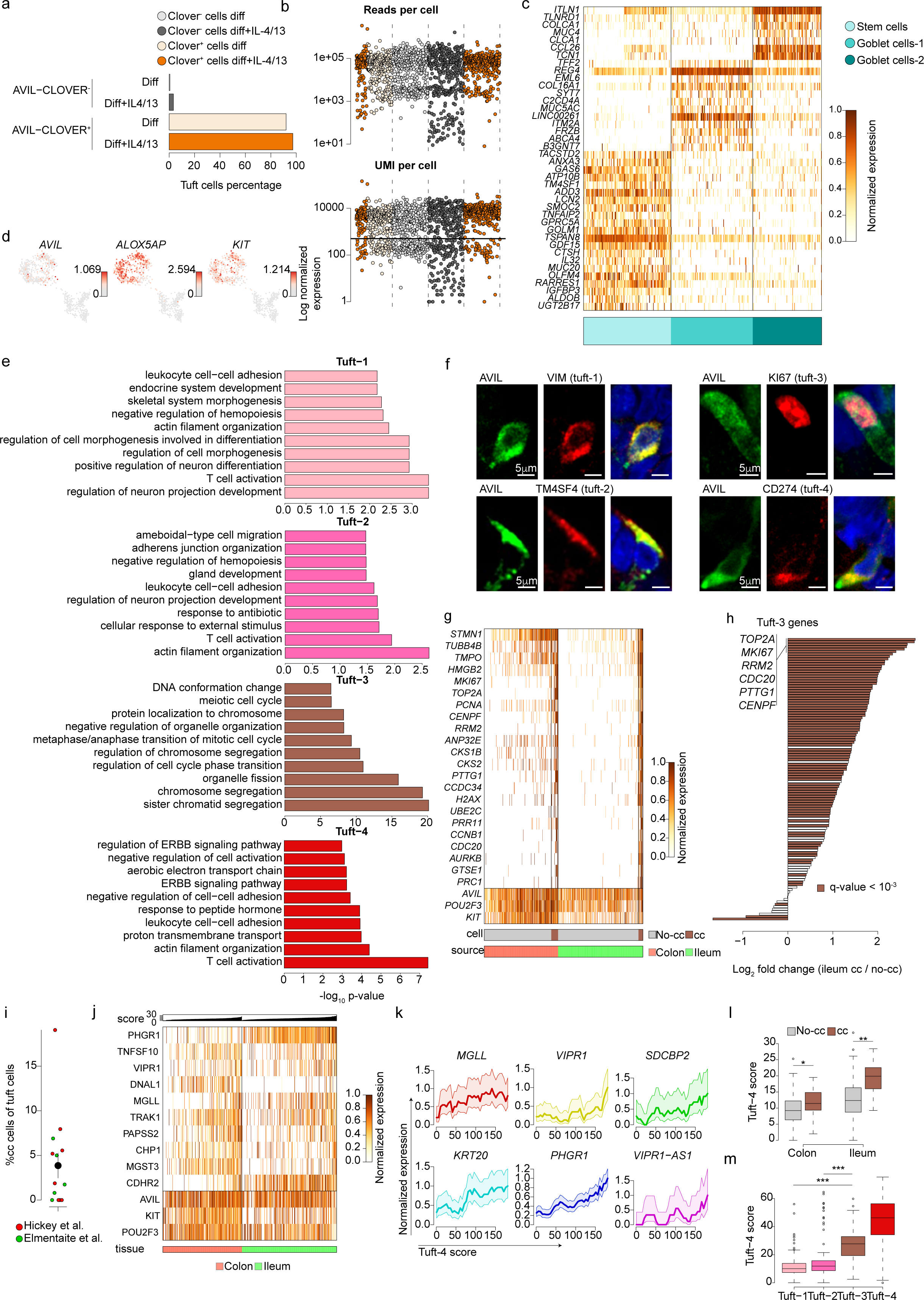
Analysis of single cell RNA sequence used in this study. a, Tuft cell frequency in different medium combinations, as determined by scRNA-seq analysis. **b,** scRNA-seq statistics, depicting number of reads per cells (top) and number of unique molecular identifiers (UMI) per cell (bottom). Each dot is a cell, colored by its gating and medium conditions. Quality threshold of 500 UMI per cell is indicated by a horizontal line. **c,** Gene expression profiles of stem cell and goblet cells. n=373 single non-tuft cells. **d,** Expression of tuft cell marker genes projected on the metacell 2D layout as in Fig. 3b-c. Dot color indicate log normalized expression. n=953 single cells. **e,** Top 10 Gene ontology (GO) terms enriched in DEG of tuft-1-4, when compared to the non-tuft cells (Fig. 3f). **f,** Representative fluorescence images of tuft-1-4 specific markers (red, as indicated) in human ileum AVIL-reporter organoids cultured in tuft cell differentiation medium (tuft 1-2) or supplemented with IL-4/IL-13 (tuft 3-4). n = 2 independent experiments on donor 1. **g**, Gene expression profiles of primary KIT^+^ tuft cells from human ileum and colon tissue. Shown are tuft-3 genes, as well as *KIT*, *POU2F3* and *AVIL*. Cells are colored by their tissue origin, and by classification into cycling cells (cc) or non-cycling cells (no-cc). n= 271 single colon and 311 single ileum KIT^+^ cells. **h**, Differential expression of all tuft-3 genes between primary ileal cycling and non-cycling KIT^+^ cells as in **g**. Genes with significant differential expression (ξ2 test; FDR-adjusted p-value < 10^-3^) are colored. **i**, Estimation of the fraction of proliferating tuft cells across 14 human primary intestine tissues from two published scRNA-seq datasets^18,25^. **j**, Gene expression profiles of primary KIT^+^ non-cycling tuft cells as in **g**. Shown are tuft-4 genes, as well as *KIT*, *POU2F3* and *AVIL*. Cells are ordered by their expression of the aggregated tuft-4 program, and colored by their tissue origin. **k**, Expression patterns of six genes along the tuft-4 activation gradient in KIT^+^ non-cycling ileal cells. Shadings indicate 95% confidence in binomial estimation of the mean. **l**, Aggregate expression of the tuft-4 program in cycling and non-cycling cells in colon and ileum. **m**, Aggregate expression of the tuft-4 program across tuft cell substates in organoids as in Fig. 3g. n=573 single tuft cells from organoids. **l-m**, Two sided Mann-Whitney test; box plots present the third quartile (top of the box), median (center lines) and first quartile (bottom of the box) of measurements. The whiskers represent 1.5 times the interquartile range from the top or bottom of the box. Diff: human tuft cell differentiation medium; cc: cell cycle. * P < 0.05, ** P < 0.01, *** P < 0.001.

**Extended Data Fig. 5.**
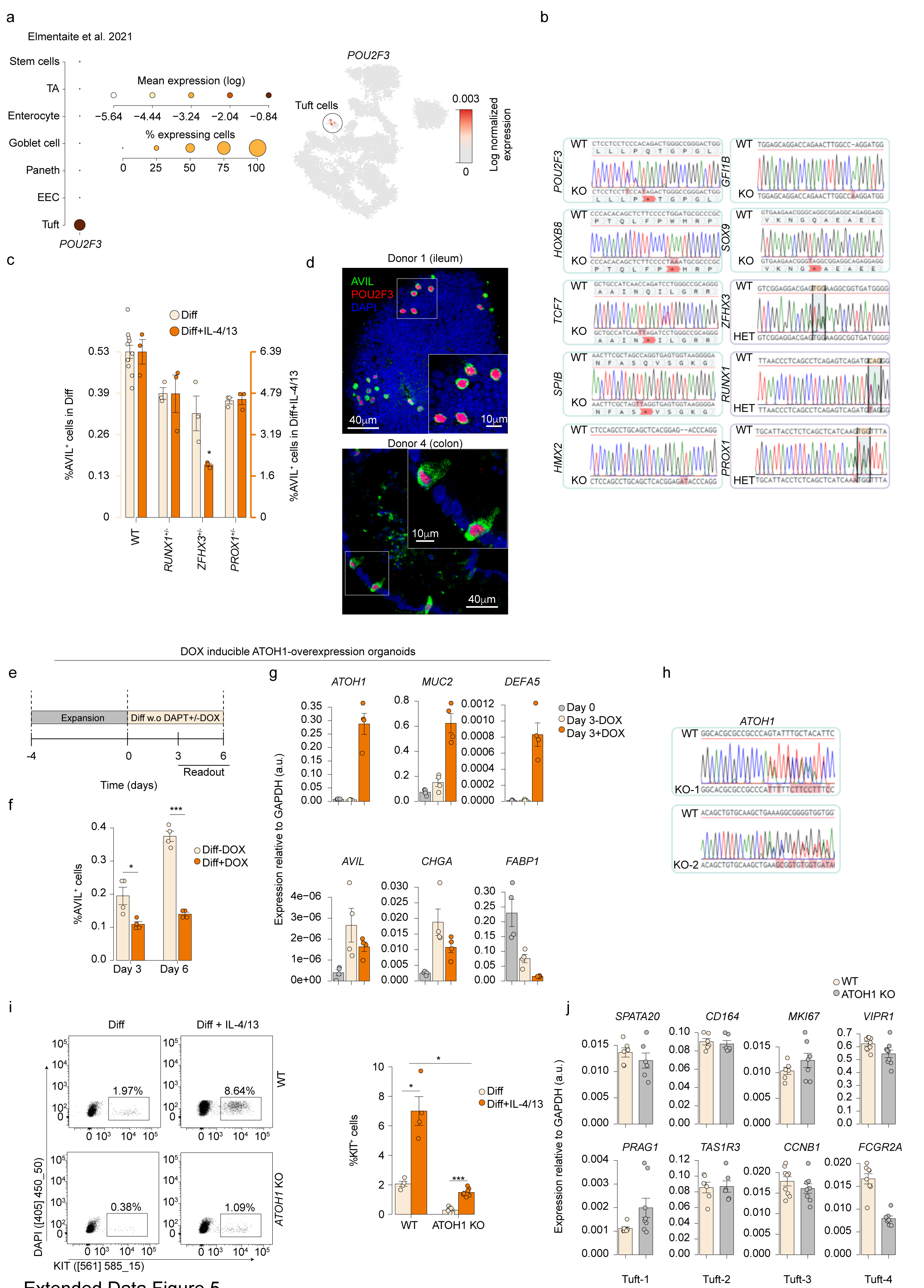
Transcription factor knock outs and overexpression in human ileum organoids. **a,** Expression of *POU2F3* across human healthy adult intestine tissue scRNA-seq dataset as in 1a. Left: Dot color relates to mean expression values and dot size to fraction of expressing cells. Right: dot color indicates log normalized expression. **b,** Genotypes of clonal transcriptional factor knock outs generated from human ileum AVIL-Clover reporter organoids. Homozygous knock outs of *POU2F3*, *HOXB8*, *TCF7*, *SPIB*, *SOX9* and heterozygous knock outs of *ZFHX3*, *RUNX1*, *PROX1* are generated using base editing (C to T) technology to induce stop codon (TAG, TAA) within exons. *HMX2*^-/-^ and *GFI1B*^-/-^ lines are generated by using conventional CRISPR-Cas9 method to induce frameshift. **c,** Quantification of AVIL^+^ cell frequency in heterozygous knock out organoid lines by flow cytometry. Each dot is one well. One of 3 independent experiments is shown, see Supplementary Information Fig. 3. WT measurements are pooled from 4 experiments. Error bars indicate SE. FDR-adjusted two-sided Student’s t-test against the WT levels. **d,** Fluorescence images of differentiated AVIL-Clover organoids, depicting AVIL (green), POU2F3 (red) and DAPI (blue) from 2 donors (donors 1, 4). **e,** Schematics of the experimental set-up for **(f-g)**. ATOH1-inducible human ileum organoids were expanded for 4 days, then differentiated in tuft cell medium without DAPT, with or without a doxycycline pulse. **f,** Quantification of AVIL^+^ cell frequency in ATOH1-inducible organoids (as in **e**) by flow cytometry. Each dot is a well. One of 3 independent experiments on donor 1 was shown, see Supplementary Information Fig. 3. Error bars indicate SE. Two-sided Student’s t-test. **g,** qPCR quantification of *ATOH1* expression and intestinal epithelial lineage markers. Each dot is a well. One of 2 independent experiments was shown, Supplementary Information Fig. 3. **h,** Genotype of clonal *ATOH1* knock outs generated from human ileum organoids. **i,** Representative flow cytometric analysis (left) and quantification (right) of KIT^+^ cell frequency in *ATOH1* knock out organoids. Organoids were differentiated for 7 days in depicted media. Each dot is a well. Results are pooled from two *ATOH1* knockout clonal lines from donor 1 (for 3 additional experiments, see Supplementary Information Fig. 3). Error bars indicate SE. Two-sided Student’s t-test. **j,** qPCR quantification of tuft-1-4 characteristic genes in KIT^+^ cells sorted from WT and *ATOH1*-knockout organoids. Organoids were differentiated for 7 days in tuft cell differentiation medium with IL-4/IL-13- removal of DAPT. Each dot is a technical replicate. n= 3 wells pooled from two *ATOH1* knockout lines. Error bars indicate SE. Diff: human tuft cell differentiation medium; TA: Transit-Amplifying Cells; EEC: Enteroendocrine cells; WT: wildtype; SE: standard error. *P < 0.05; **P< 0.01; ***P< 0.001.

**Extended Data Fig. 6.**
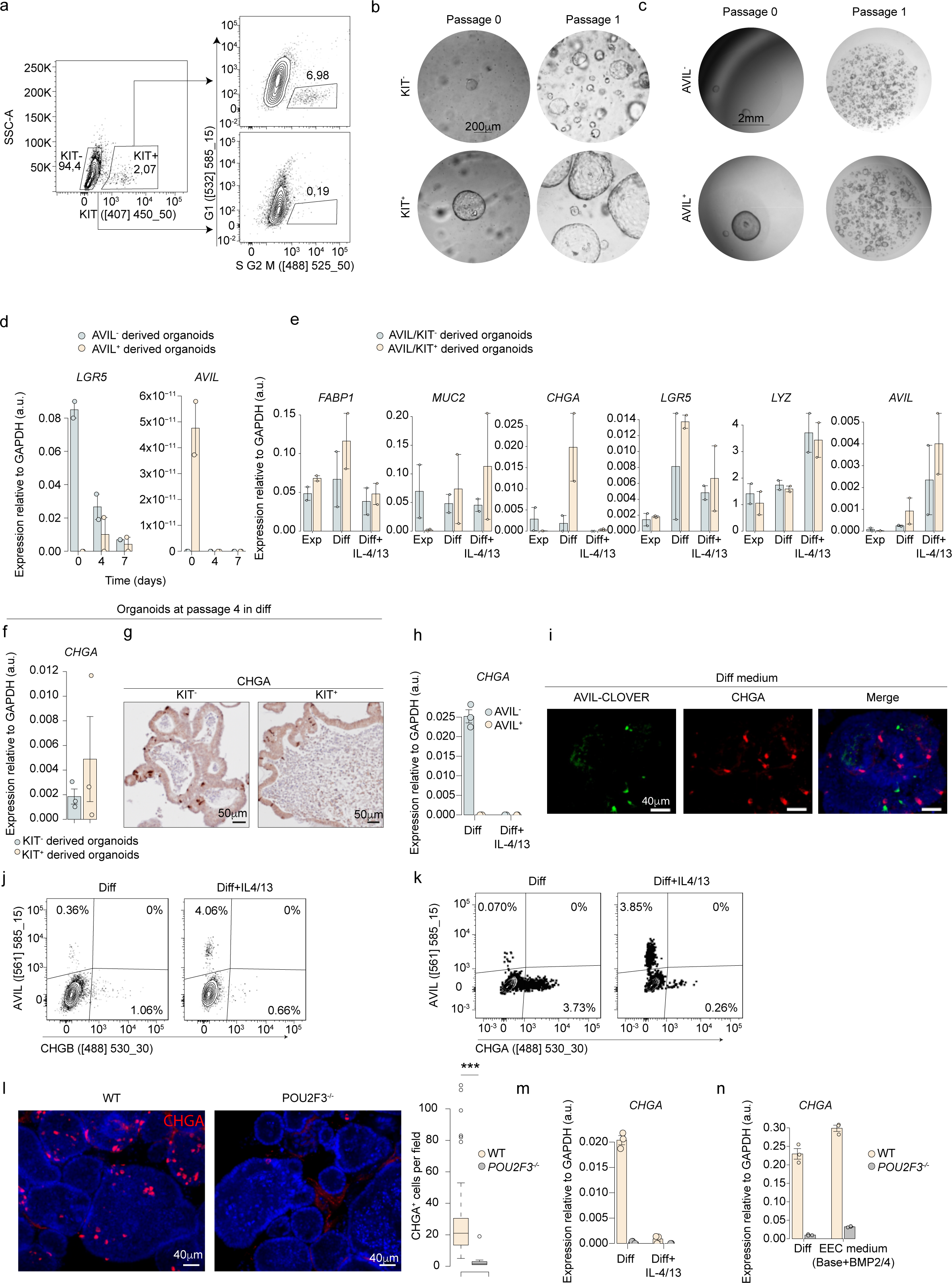
Human intestinal tuft cell derived organoids. **a,** Gating strategy for organoid outgrowth from single S/G2/M phase tuft cell (KIT^+^) and non-tuft cell (KIT^-^) in FUCCI reporter organoids. Prior to sorting, organoids were differentiated for 3 days in tuft cell medium with IL-4 and IL-13. **b-c,** Representative images of organoid outgrowth from single KIT^+^ and KIT^-^ cells in human adult duodenum tissue **(b)**, and from AVIL^+^ and AVIL^-^ cells in AVIL-Clover reporter organoids **(c)**. **(b)** n= 3 donors, Supplementary Information Fig. 4. **(c)** n= 2 donors; donor 1 is shown, donor 2 see Supplementary Information Fig. 4. **d,** qPCR quantification of *LGR5* and *AVIL* expression in sorted AVIL^+^ and AVIL^-^ cells at depicted time points following seeding. Sorted cells were cultured in human expansion medium. Each dot is one donor (donors 1-2). Error bars indicate SE. **e,** qPCR quantification of lineage markers representing different intestinal epithelial cell types in organoids after first passage. Each dot is one donor (donors 1-2). Error bars indicate SE. **f-g,** qPCR quantification **(f)** and visualization **(g)** of CHGA expression in organoids of passage 4 derived from KIT^+^ and KIT^-^ cells as in (a). Each dot is a well. 3 independent experiments were performed on two donors (donor 1 is shown; for donor 2, see Supplementary Information Fig. 4). Error bars indicate SE. **h**, qPCR quantification of *CHGA* expression in AVIL^+^ and AVIL^-^ cells sorted from AVIL-Clover reporter organoids. Each dot is a well. Results are pooled from 3 independent experiments on donor 1. Error bars indicate SE. **i,** Representative fluorescence image of an AVIL-Clover (green) differentiated organoid co-stained with CHGA (red), and DAPI (blue). n= 3 independent experiments on donor 1. **j-k,** Representative flow cytometric analysis of AVIL and CHGB **(h)** or CHGA **(i)** in human ileum AVIL-P2A-tdtomato/CHGB-mNeon **(j)** and AVIL-P2A-tdtomato/CHGA-Clover **(k)** double reporter organoid lines. Organoids were differentiated in tuft cell medium with or without IL- 4/IL-13 for 4 days **(j)** or 9 days **(k)**. n= 3 independent experiments on donor 1. **l,** Representative fluorescence images (left) and quantification (right) of CHGA^+^ cells in WT and *POU2F3*^-/-^ organoids differentiated for 7 days in tuft cell differentiation medium. Results are pooled from 2 independent experiments on donor 1, n = 76 (WT) and 46 (*POU2F3*^-/-^) individual organoids. Two-sided Student’s t-test. The box plot presents the third quartile (top of the box), median (center lines) and first quartile (bottom of the box) of measurements. The whiskers represent 1.5 times the interquartile range from the top or bottom of the box. **m,** qPCR quantification of *CHGA* expression in WT and *POU2F3^-/-^* organoids differentiated in tuft cell differentiation medium with or without IL-4 and IL-13. Each dot is a well. Experiments were performed on two donors (donor 1 is shown; for donor 4, see Supplementary Information Fig. 4). Error bars indicate SE. **n,** Same as **(m)**, organoids were differentiated in either tuft cell or EEC differentiation medium. Each dot is a well. Results are pooled from 3 independent experiments on donor 1. Error bars indicate SE. WT: wildtype; Diff: human tuft cell differentiation medium; SE: Standard error. ***P< 0.001.

**Extended Data Fig. 7.**
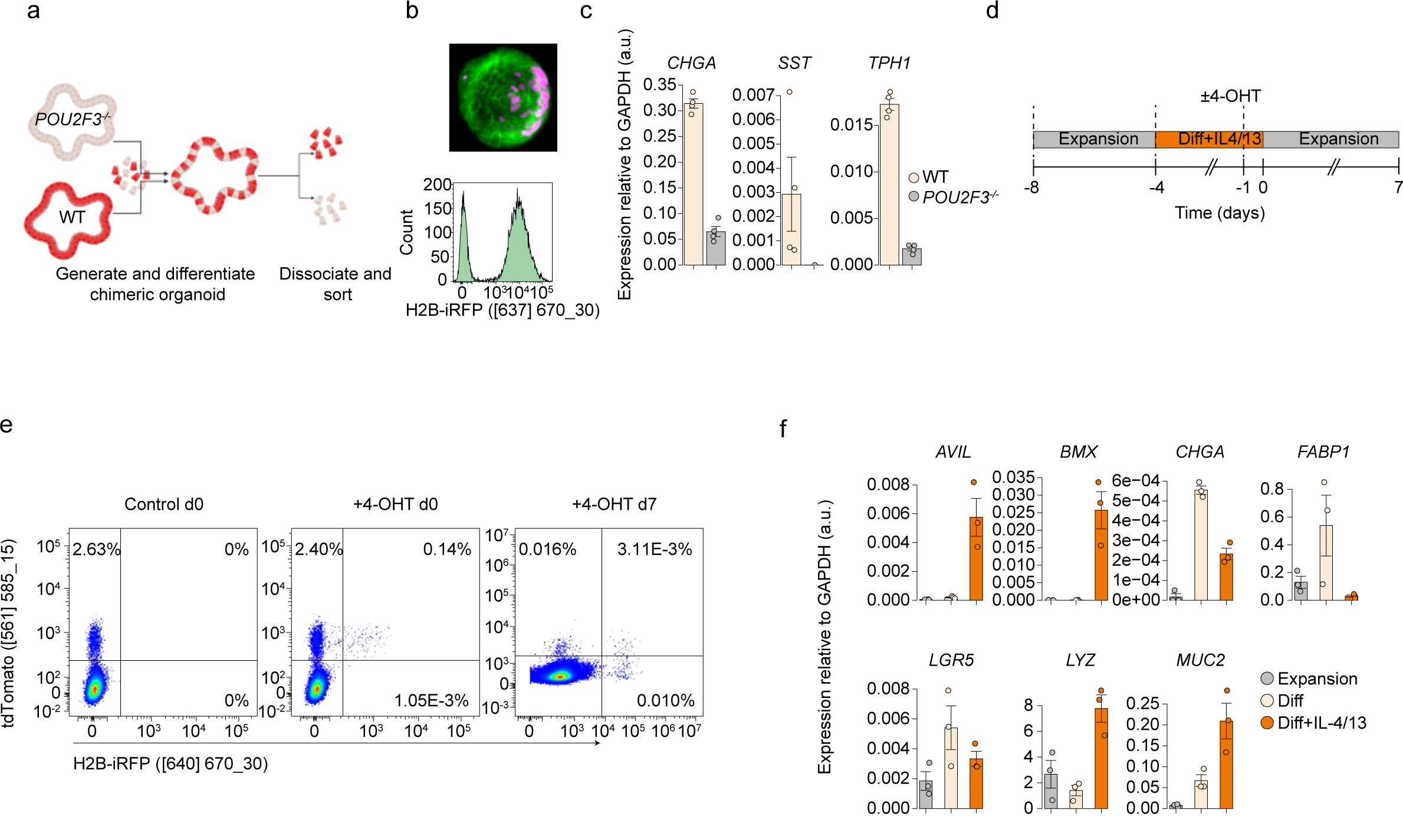
Lineage tracing of human AVIL^+^ tuft cells. **a,** Schematics of experimental set-up for **(b-c)**. Mosaic organoids, derived from *POU2F3^-/-^* and H2B-iRFP wildtype organoid lines, were differentiated for 7 days in tuft cell differentiation medium before sorting. **b**, Representative fluorescence image (top) and flow cytometric analysis (bottom) of mosaic organoids as in **(a)**. **c**, qPCR analysis (right) showing expression of enteroendocrine cell markers (*CHGA, SST, TPH1*) from sorted H2B-iRFP^+^ and iRFP^-^ (i.e., *POU2F3^-/-^)* cells. Each dot is a well. Results are pooled from 2 independent experiments on donor 1. Error bars indicate SE. **d, e,** Schematics of experimental set-up **(d)** and representative flow cytometric analysis **(e)** of AVIL lineage tracing organoids. Organoids were differentiated for 4 days in tuft cell medium with IL-4/IL-13, with or without exposure to 1 μM 4-Hydroxytamoxifen for 20 hours, then medium was changed to human intestinal expansion medium. n = 3 independent experiments. **f,** qPCR quantification of intestinal epithelial lineage markers in traced organoids derived from single AVIL^+^iRFP^+^ cells sorted at day 0 as in **(d)**. Each dot is a well. Results are pooled from 3 independent experiments on donor 1. Error bars indicate SE. Diff: human tuft cell differentiation medium; WT: wildtype; SE: standard error.

**Extended Data Fig. 8.**
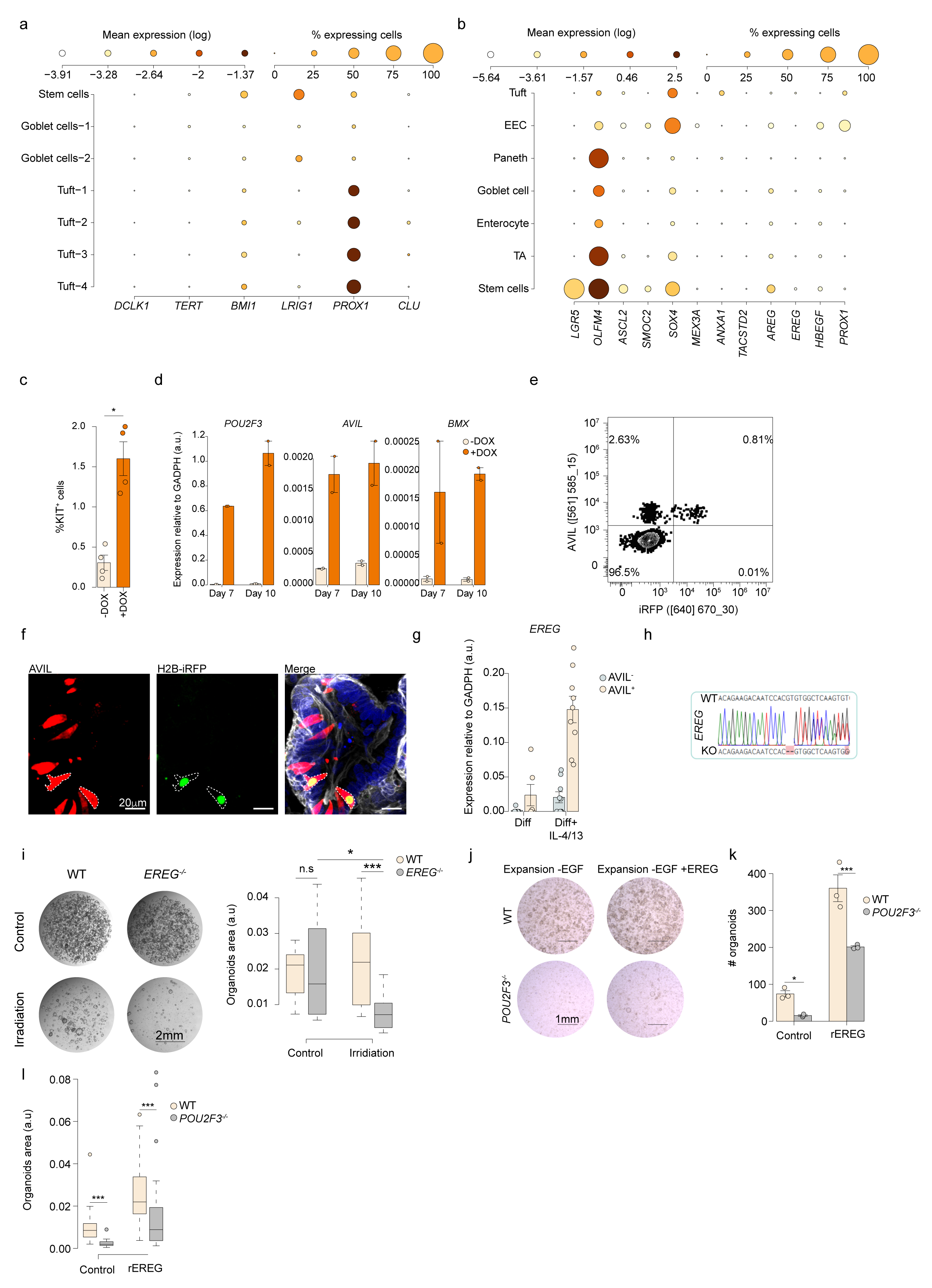
Regeneration potential of tuft cells following damage. **a-b,** Expression of selected genes associated with regenerative stem cells across ileum organoid-derived subsets **(a)**, and across primary epithelial cells as in 1a **(b)**. Dot color relates to mean expression values and dot size to fraction of expressing cells. n=953 **(a)** and 15,184 **(b)** single cells. **c-d,** Quantification of KIT^+^ cell frequency **(c)** and qPCR quantification of tuft cell genes **(d)** in DOX-triggered POU2F3 overexpression organoids. Organoids were differentiated in tuft cell medium. One of 2 **(d)** or 3 **(c)** independent experiments on donor 2 is shown (Supplementary Information Fig. 5). Error bars indicate SE. Two-sided Student’s t-test. **e–f,** Representative flow cytometric analysis **(e)** and fluorescence image (**f)** of AVIL- lineage tracing organoids after irradiation. n= 3 independent experiments on donor 1. **g,** qPCR quantification of *EREG* expression in sorted AVIL^+^ and AVIL^-^ cells from human ileum organoids. Organoids were differentiated in tuft cell differentiation medium with and without IL-4 and IL-13 for 7 days. Each dot represents a technical replicate, results is pooled from 2 or 3 wells of donor 1, also see Supplementary Information Fig. 5. Error bars indicate SE. **h,** Genotype of human ileum *EREG* knock out organoids. **i,** Representative images (left) and quantification of organoid area (right) from WT and *EREG*^-/-^ organoids exposed to IL-4 and IL-13 after irradiation (as Fig 5e). Results are pooled from 2 independent experiments on donor 1, n= 600-900 individual organoids. Two-sided Student’s t-test. **j–l,** Wildtype and *POU2F3^-/-^* organoids were differentiated for 7 days in tuft cell differentiation medium with IL-4 and IL-13, passaged, then cultured for 7 days in human intestinal expansion medium by removal of EGF, with or without recombinant EREG (rEREG). Shown are representative images **(j)**, quantification of organoid numbers **(k)**, and organoid areas **(l)**. Three independent experiments were performed on 2 donors. (donor 1 is shown; for donor 5, see Supplementary Information Fig. 5). **k,** Each dot is a well. Error bars indicate SE. Two-tailed Mann-Whitney test. **l,** Each dot is an individual organoid, n = 20-40 organoids per condition. Two-sided Student’s t-test. **i,l,** box plots present the third quartile (top of the box), median (center lines) and first quartile (bottom of the box) of measurements. The whiskers represent 1.5 times the interquartile range from the top or bottom of the box. Diff: human tuft cell differentiation medium; WT: wildtype; TA: Transit-Amplifying Cells; EEC: Enteroendocrine cells; SE: standard error; n.s: not significant. *P < 0.05, ***P< 0.001.

**Extended Data Fig. 9.**
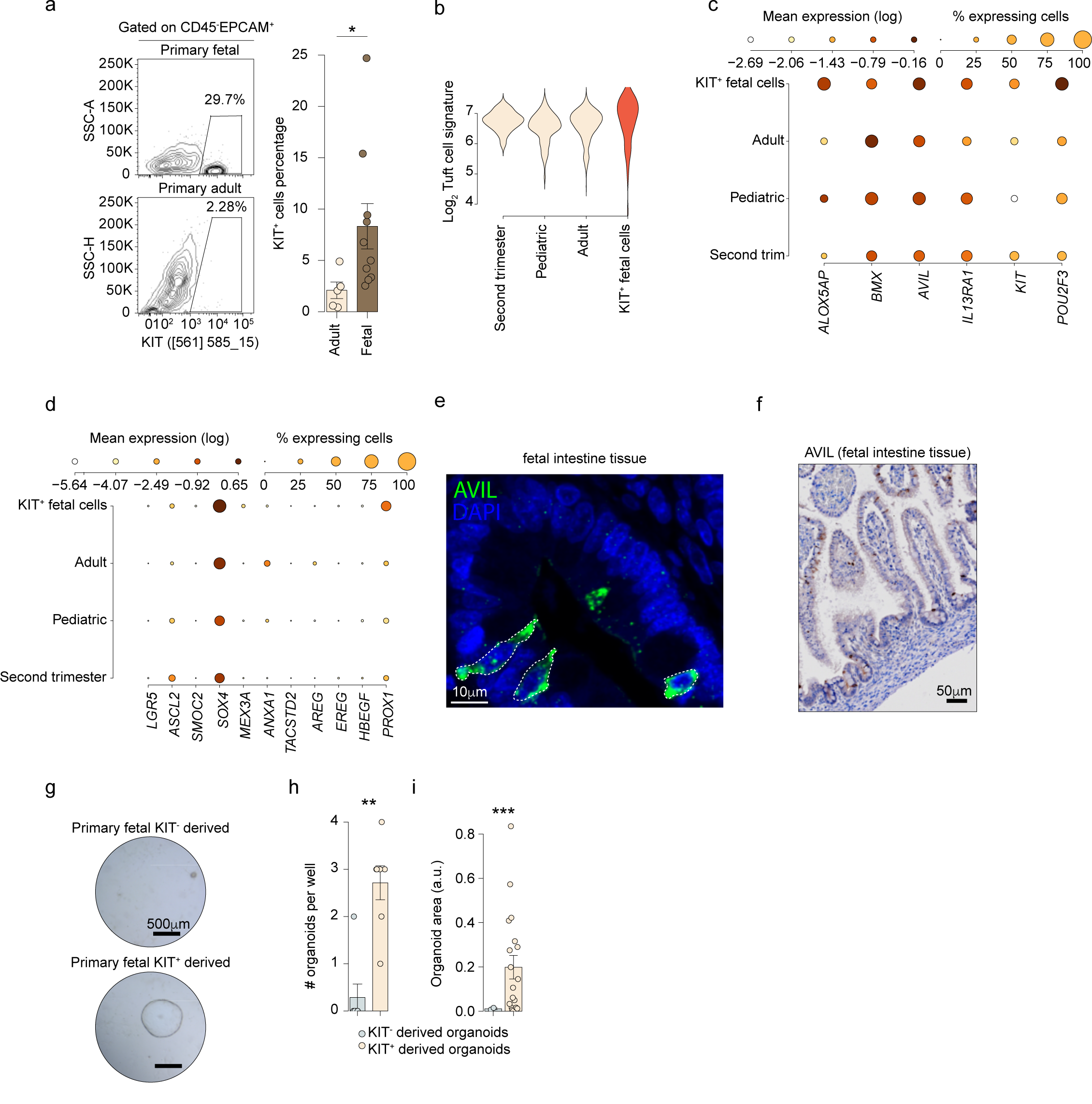
Analysis of human fetal intestinal tuft cells. **a,** Representative flow cytometric analysis (left) and quantification (right) of KIT^+^ cells frequency of human fetal and adult intestinal tissue. Single cells are pre-gated on DAPI^-^ CD45^-^ EPCAM^+^. Each dot is a donor. Error bars indicate SE. Two-sided Student’s t-test. **b,** Distribution of a tuft cell signature across human tuft cells from different development stages^18^, as well as within a sorted population of fetal KIT^+^ cells. The tuft cell signature is as in fig. 2h. **c-d,** Expression of core tuft cells genes **(c)** and genes associated with regenerative stem cells **(d)** across human tuft cells from different development stages, as well as in a sorted population of fetal KIT^+^ cells. Dot color relates to mean expression values and dot size to fraction of expressing cells. **b-d,** n=699 tuft cells, and 89 KIT^+^ fetal cells. **e-f,** Representative images of AVIL expression in histological sections of human fetal intestine. n= 3 donors. **g-i,** Representative images **(g)** and quantifications of organoid numbers **(i)** and area **(j)** of organoids derived from single primary fetal (week 19-21) KIT^+^ and KIT^-^ cells. One of three donors is shown, see Supplementary Information Fig. 5. Error bars indicate SE. **h**, Each dot is a well; two-tailed Mann-Whitney test. **i**, Each dot is an individual organoid; two-sided Student’s t-test. SE: Standard error. *P < 0.05, **P<0.01, ***P< 0.001.

